# The cryptic plastid of *Euglena longa* defines a new type of non-photosynthetic plastid organelles

**DOI:** 10.1101/765255

**Authors:** Zoltán Füssy, Kristína Záhonová, Aleš Tomčala, Juraj Krajčovič, Vyacheslav Yurchenko, Miroslav Oborník, Marek Eliáš

## Abstract

Most secondarily non-photosynthetic eukaryotes have retained residual plastids whose physiological role is often still unknown. One such example is *Euglena longa,* a close non-photosynthetic relative of *Euglena gracilis* harbouring a plastid organelle of enigmatic function. By mining transcriptome data from *E. longa* we finally provide an overview of metabolic processes localized to its elusive plastid. The organelle plays no role in biosynthesis of isoprenoid precursors and fatty acids, and has a very limited repertoire of pathways concerning nitrogen-containing metabolites. In contrast, the synthesis of phospholipids and glycolipids has been preserved, curiously with the last step of sulfoquinovosyldiacylglycerol synthesis being catalysed by the SqdX form of the enzyme so far known only from bacteria. Notably, we show that the *E. longa* plastid synthesizes tocopherols and a phylloquinone derivative, the first such report for non-photosynthetic plastids studied so far. The most striking attribute of the organelle is the presence of a linearized Calvin-Benson (CB) pathway including RuBisCO yet lacking the gluconeogenetic part of the standard cycle, together with ferredoxin-NADP^+^ reductase (FNR) and the ferredoxin/thioredoxin systems. We hypothesize that FNR passes electrons to the ferredoxin/thioredoxin systems from NADPH to activate the linear CB pathway in response to the redox status of the *E. longa* cell. In effect, the pathway may function as a redox valve bypassing the glycolytic oxidation of glyceraldehyde-3-phosphate to 3-phosphoglycerate. Altogether, the *E. longa* plastid defines a new class of relic plastids that is drastically different from the best studied organelle of this category, the apicoplast.

**Importance:** Colourless plastids incapable of photosynthesis evolved in many plant and algal groups, but what functions they perform is still unknown in many cases. Here we study the elusive plastid of *Euglena longa,* a non-photosynthetic cousin of the familiar green flagellate *Euglena gracilis.* We document an unprecedented combination of metabolic functions that the *E. longa* plastid exhibits in comparison with previously characterized non-photosynthetic plastids. For example, and truly surprisingly, it has retained the synthesis of tocopherols (vitamin E) and a phylloquinone (vitamin K) derivative. In addition, we offer a possible solution of the long-standing conundrum of the presence of the CO_2_-fixing enzyme RuBisCO in *E. longa.* Our work provides a detailed account on a unique variant of relic plastids, the first among non-photosynthetic plastids that evolved by secondary endosymbiosis from a green algal ancestor, and suggests that it has persisted for reasons not previously considered in relation to non-photosynthetic plastids.

## INTRODUCTION

Photosynthesis was supposedly the primary evolutionary advantage driving the acquisition of the primary plastid as well as its further spread in eukaryotes by secondary and higher-order endosymbioses (1–3). However, plastids host many other metabolic pathways, such as biosynthesis of amino and fatty acids, isopentenyl pyrophosphate (IPP) and its derivatives (isoprenoids), and tetrapyrroles (4–6). Hence, reversion of photosynthetic lineages to heterotrophy typically does not entail plastid loss and non-photosynthetic plastids are found in many taxa (7–10).

The most extensively studied relic plastid is the apicoplast of apicomplexan parasites *(Plasmodium falciparum* and *Toxoplasma gondii,* above all). The essentiality of the apicoplast for parasite survival has attracted much attention as a promising target for parasite-specific inhibitors (11, 12). So far, three plastid pathways seem to condition the apicoplast retention: non-mevalonate IPP synthesis, haem synthesis, and type II fatty acid synthesis (FASII) (13). Less is known about plastid metabolic functions in other non-photosynthetic algal lineages. Many of them have a metabolic capacity similar to the apicoplast (10, 14, 15), but some house a more complex metabolism that includes amino acid biosynthesis and carbohydrate metabolism pathways (16–18). Until recently, IPP synthesis appeared to be a process conserved even in the most reduced plastids, such as the genome-lacking plastids of certain alveolates (8, 19). However, non-photosynthetic plastids lacking this pathway are now documented (9, 20, 21). Thus, there generally is a metabolic reason for plastid retention, although the cases of plastid dependency differ between lineages.

Like their prime representative *Euglena gracilis*, most euglenophytes are mixotrophs containing complex three-membrane-bound plastids derived from a green alga (22–24). Non-photosynthetic mutants of *E. gracilis* are capable of heterotrophic living (reviewed in 7, 25) and several euglenophyte lineages independently became secondarily heterotrophic (26). The best known is *Euglena* (previously *Astasia) longa*, a close relative of *E. gracilis* (26, 27). Although documentation at the cytological level is spurious (28–30), molecular sequence data provide clear evidence for the presence of a cryptic plastid organelle in this species. The *E. longa* plastid genome was sequenced two decades ago (31) and shown to lack all the photosynthesis-related genes, surprisingly except for *rbcL* encoding the large subunit of ribulose-1,5-bisphosphate carboxylase/oxygenase (RuBisCO). More recently, the existence of a nucleus-encoded small RuBisCO subunit (RBCS), synthesized as a precursor polyprotein, was documented in *E. longa,* although its processing into monomers could not be demonstrated (32). The physiological role of the *E. longa* RuBisCO and the whole plastid remains unknown, but indirect evidence suggests that the plastid is essential for *E. longa* survival (33–36).

To provide a resource for investigating the biology of *E. longa* and its plastid, we generated a transcriptome assembly and demonstrated its high completeness and utility (37). We also showed that nucleus-encoded plastidial proteins in *E. longa* employ an N-terminal plastid-targeting bipartite topogenic signal (BTS) of the same two characteristic classes as known from *E. gracilis.* The *E. longa* transcriptome revealed unusual features of the plastid biogenesis machinery shared with photosynthetic euglenophytes, but also suggested specific reductions of housekeeping functions, reflecting the loss of photosynthesis (37). Nevertheless, the anabolic and catabolic pathways localized to the *E. longa* colourless plastid have not been characterized. Hence, we set to exploit the available sequence data to chart the metabolic map of the *E. longa* plastids. The analyses were greatly facilitated by the recent characterization of the *E. gracilis* plastid metabolic network based on a proteomic analysis of the organelle (38). Our study provides the first comprehensive view of a non-photosynthetic secondary plastid of green-algal origin and shows that the metabolic capacity of the *E. longa* plastid is strikingly different from those of the apicoplast and other relic plastids characterized in sufficient detail.

## RESULTS

### The plastid protein complement of *E. longa* is dramatically reduced compared to that its photosynthetic cousin

To obtain a global view of the repertoire of the plastid proteins in *E. longa,* we searched its transcriptome assembly to identify putative orthologs of the proteins defined as part of the *E. gracilis* plastid proteome (38). Of the 1,312 such proteins encoded by the *E. gracilis* nuclear genome, less than a half – namely 594 – exhibited an *E. longa* transcript that met our criteria for orthology (Table S1). As expected, the functional categories with the least proportion of putative *E. longa* orthologs included “photosynthesis”, “metabolism of cofactors and vitamins”, and “reaction to oxidative and toxic stress” with 95.89%, 85.11%, and 73.33% of proteins missing in *E. longa.* Interestingly, *E. longa* lacks counterparts also of some plastidial proteins involved in gene expression or genome maintenance, suggesting that the metabolic simplification, primarily the loss of photosynthesis itself with its high demand on protein turnover and mutagenic effects on the plastid genome, may have relaxed the constraints on the respective house-keeping molecular machineries.

Although these results clearly demonstrate dramatic reduction of the functional complexity of the *E. longa* plastid when compared to the plastid of its photosynthetic relative, they should not be interpreted such that the plastid harbours exactly the ~600 proteins identified by the orthology search. Firstly, the proteomically defined set of the putative *E. gracilis* plastid proteins is certainly affected by the presence of false negatives (bona fide plastid proteins missed by the analysis) as well as false positives (contaminants; (38). Secondly, orthology does not necessarily imply the same subcellular localization. Hence, to obtain a finer view of the physiological functions of the *E. longa* plastid, we systematically searched for homologs of enzymes underpinning metabolic pathways known from plastids in general. N-terminal regions of the candidates were evaluated for characteristics of presequences predicting a specific subcellular localization to distinguish those likely representing plastid-targeted proteins from enzymes located in other compartments. Some of the bioinformatic predictions were further tested by biochemical analyses.

### *E. longa* plastid lacks the MEP pathway of IPP biosynthesis, yet has kept the production of tocopherol and a phylloquinone derivative

There are two parallel pathways of IPP biosynthesis in *E. gracilis* (39): the mevalonate (MVA) pathway localized to the mitochondrion (first three enzymes) and the cytosol (the rest), and the plastid-localized 2-C-methyl-D-erythritol (MEP) pathway, the latter providing precursors for synthesis of terpenoid compounds connected to photosynthesis, namely carotenoids and plastoquinone (38, 39). As expected, only enzymes of the MVA pathway were found in *E. longa* (Table S2, Fig. 1a). The carotenoid and plastoquinone biosynthesis enzymes are all missing, but surprisingly the *E. longa* plastid appears still involved in terpenoid metabolism, specifically in its phytol branch.

**Figure 1:**
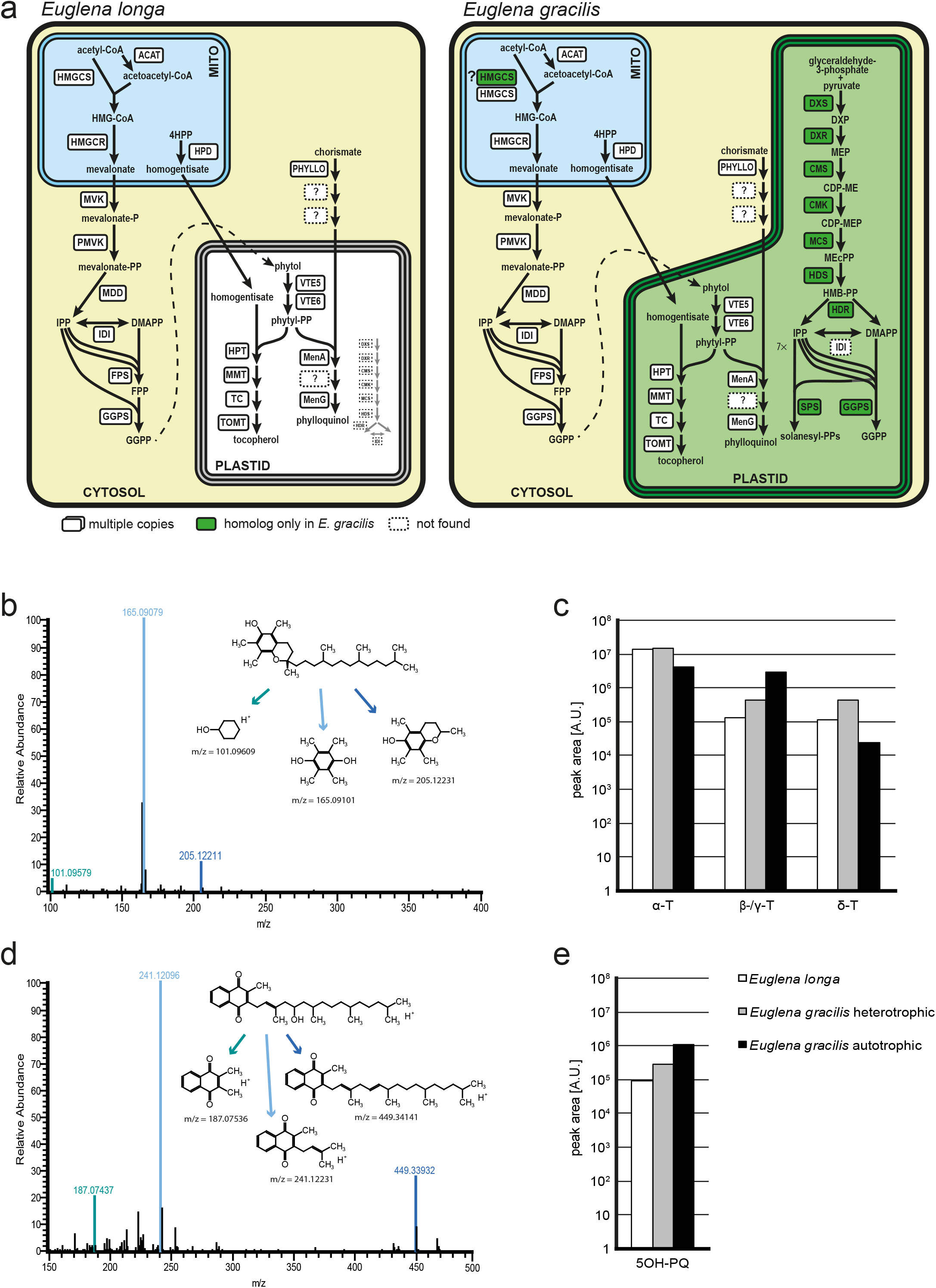
IPP and terpenoid-quinone biosynthesis in *E. longa* and its phototrophic relative *E. gracilis.* **a**: Schematic comparison of the localization and evolutionary origin of enzymes (see colour-coding graphical legend below the “cells”). Abbreviations, IPP synthesis: ACAT – acetyl-CoA acetyltransferase, CDP-ME – 4-(cytidine 5’-diphospho)-2-C-methyl-D-erythritol, CDP-MEP – 2-phospho-CDP-ME, CMK – CDP-ME kinase, CMS – CDP-ME synthase, DMAPP – dimethylallyl diphosphate, DXP – 1-deoxy-D-xylulose 5-phosphate, DXR – DXP reductase, DXS – DXP synthase, FPP – farnesyl siphosphate synthase, GGPS – geranylgeranyl-diphosphate synthase, HDR – HMB-PP reductase, HDS – HMB-PP synthase, HMB-PP – 4-hydroxy-3-methylbut-2-en-1-yl diphosphate, HMG-CoA – 3-hydroxy-3-methylglutaryl-CoA, HMGCR – HMG-CoA reductase, HMGCS – HMG-CoA synthase, IDI – isopentenyl-diphosphate delta-isomerase, MCS – MEcPP synthase, MDD – mevalonate-diphosphate decarboxylase, MEcPP – 2-C-methyl-D-erythritol 2,4-cyclodiphosphate, MEP – 2-C-methyl-D-erythritol 4-phosphate, MVK – mevalonate kinase, PMVK – phosphomevalonate kinase, PPS – unspecified polyprenyl-diphosphate synthase, ? – unclear substrate; Terpenoid-quinone synthesis: 4HPP – 4-hydroxyphenylpyruvate, HPD – hydroxyphenylpyruvate dioxygenase, HPT – homogentisate phytyltransferase, MMT – MPBQ/MPSQ methyltransferase, TAT – tyrosine aminotransferase, TC – tocopherol cyclase, TMT – tocopherol-O-methyltransferase, VTE5 – phytyl kinase, VTE6 – phytyl-phosphate kinase. **b**: MS/MS spectrum record of *E. longa* α-tocopherol and the proposed fragmentation pattern in positive ionization mode (inset). Monoisotopic masses of particular fragments were obtained by simulation in Xcalibur software. c: Semiquantitative comparison of tocopherol species in *E. longa,* heterotrophically (dark) grown *E. gracilis* and autotrophically grown *E. gracilis.* **d-e**: MS/MS spectrum record of *E. longa* 5-hydroxyphylloquinone and the proposed fragmentation pattern in positive ionization mode (inset); semiquantitative comparison of 5-hydroxyphylloquinone in *E. longa,* heterotrophically (dark) grown *E. gracilis* and autotrophically grown *E. gracilis*.

Photosynthetic eukaryotes generally produce three types of phytol derivatives, tocopherols (vitamin E), phylloquinone (PhQ; vitamin K1) and chlorophyll, starting with a common precursor phytyl-PP, which is (directly or indirectly *via* salvage of phytol liberated by chlorophyll degradation) made by reduction of geranylgeranyl-PP derived from the MEP pathway (40). *E. gracilis* proved to be unusual not only because it lacks the conventional geranylgeranyl-PP reductase (38), but also for making phytol from a precursor provided by the MVA pathway (39, 41). The route of phytol synthesis is currently unknown, though phytyl-PP might be synthesized in the *E. gracilis* plastid exclusively by the step-wise phosphorylation of phytol by phytol kinase (VTE5) and phytyl phosphate kinase (VTE6), enzymes employed in phytol salvage (38). *E. longa* has retained both VTE5 and VTE6, each being highly similar to their *E. gracilis* orthologs and exhibiting putative BTS (Fig. S1; Table S2). Since *E. longa* lacks chlorophyll and hence phytol recycling, these two enzymes are likely to participate in the *de novo* synthesis of phytol.

*E. gracilis* is known to make tocopherols and a PhQ derivative, 5’-monohydroxyphylloquinone (OH-PhQ; 38, 42, 43). All four enzymes mediating synthesis of α-tocopherol from phytyl-PP and homogentisate were identified and are localized to its plastid (38). Interestingly, their orthologs are found in *E. longa,* all with a typical BTS or at least with the N-terminal region being highly similar to the *E. gracilis* counterpart (Table S2), consistent with their presumed plastidial localization (Fig. 1a). Homogentisate itself is apparently made outside the plastid, as the enzyme responsible for its synthesis (4-hydroxyphenylpyruvate dioxygenase) is not found in the *E. gracilis* plastid proteome and the respective proteins have a predicted mitochondrial transit peptide in both *E. gracilis* and *E. longa* (Table S2). To test the predicted ability of *E. longa* to produce α-tocopherol, we used HPLC-MS/MS to analyse extracts from this species and *E. gracilis* (grown at two different conditions – in light and in darkness) for comparison. Tocopherols were detected in both species (Fig. 1b), with α-tocopherol being the dominant form present in equivalent amounts in all three samples (Fig. 1c). The signals of β− and/or γ-tocopherol (indistinguishable by the method employed) and of δ-tocopherol suggest that tocopherol cyclase, and possibly also tocopherol O-methyltransferase, of both *Euglena* species can process substrates with or without the 3-methyl group on the benzene ring (Fig. S2).

The synthesis of OH-PhQ in *E. gracilis* is understood only partially, with only three enzymes of the pathway previously identified at the molecular level: the large multifunctional protein PHYLLO, apparently localized to the cytosol and catalysing the first four steps leading to o-succinylbenzoate; MenA, catalysing phytylation of dihydroxynaphthoate localized in the plastid; and MenG (demethylnaphthoquinone methyltransferase), possessing a typical BTS but not directly confirmed as plastidial by proteomics (38). Strikingly, *E. longa* expresses orthologs of these three *E. gracilis* proteins, all with the same predicted subcellular localization (Fig. 1a, Table S2). Like in *E. gracilis,* no candidates for other enzymes required for OH-PhQ synthesis could be identified by homology searches in *E. longa.* Still, OH-PhQ could be detected in this species (Fig. 1d, Fig. S3), although with a significantly lower abundance compared to that in *E. gracilis* (Fig. 1e).

### *E. longa* plastid plays a limited role in the metabolism of nitrogen-containing compounds

Some of the apparent peculiarities of the *E. longa* plastid do not stem from the loss of photosynthesis, but rather reflect unusual features of the plastid in euglenophytes in general. This particularly concerns plastid functions in the metabolism of nitrogen-containing compounds. Plastids are commonly involved in nitrogen assimilation due to housing nitrite reductase (44, 45), but *E. gracilis* cannot assimilate nitrate or nitrite (46, 47). Accordingly, no nitrite reductase can be identified in the transcriptome data from this species and *E. longa.* The plastids of both *Euglena* species apparently also lack the enzymes working immediately downstream of nitrite reductase, i.e. glutamine synthetase and glutamine oxoglutarate aminotransferase (the GS/GOGAT system common in plastids of other groups; 48, 49), indicating that the plastids rely on the import of organic nitrogen, similarly to what has been recently proposed for chromerids (50) and chrysophytes (20, 21).

A surprising feature of the *E. gracilis* plastid metabolism is the paucity of amino acid-related pathways (38). *E. longa* is even more extreme in this regard, because it lacks counterparts of the plastid-targeted serine biosynthesis enzymes. Thus, we could localize only two elements of amino acid biosynthesis pathways to the *E. longa* plastid (Fig. S4): serine/glycine hydroxymethyltransferase, whose apparent role is to provide the one-carbon moiety for formylmethionyl-tRNA synthesis required for the plastidial translation; and one of the multiple isoforms of cysteine synthase A, which (like in *E. gracilis)* apparently relies on O-acetyl-L-serine synthesized outside of the plastid (see (38), and Table S3). This is not due to incompleteness of the sequence data, as the E. *longa* transcriptome encodes enzymes for the synthesis of all 20 proteinogenic amino acids, yet their predicted localization lies outside the plastid (Table S3).

Amino acids also serve as precursors or nitrogen donors for the synthesis of various other compounds in plastids (51, 52). This includes tetrapyrrole synthesis, which in *E. gracilis* is mediated by two parallel pathways localized to the mitochondrion/cytoplasm and the plastid (53). As described in detail elsewhere (Füssy, Záhonová, Oborník & Eliáš, unpublished), *E. longa* possesses the full mitochondrial-cytoplasmic pathway, whereas the plastidial one is restricted to its middle part potentially serving for synthesis of sirohaem, but not haem and chlorophyll (Fig. S4). The spectrum of reactions related to the metabolism of other nitrogen-containing cofactors or their precursors is very limited in the plastids of both *Euglena* spp. (Table S4). We identified only one such candidate in *E. longa* – vitamin B6 salvage catalysed by pyridoxamine 5’-phosphate oxidase, whereas *E. gracilis* additionally expresses two plastid-targeted isoforms of pyridoxine 4-dehydrogenase. Enzymes of *de novo* synthesis or salvage of purines and pyrimidines are also absent from the plastid of both *Euglena* species, except for a plastidial CTP synthase isoform in *E. gracilis* (supported by proteomic data), which is not expressed by *E. longa.* The lack of *in situ* CTP production may reflect the presumably less extensive synthesis of RNA and/or CDP-diacylglycerol (a precursor of phospholipids) in the *E. longa* plastid. Finally, *E. longa* expresses an ortholog of spermidine synthase found in the plastid proteome of *E. gracilis*, but it has a modified N-terminal sequence not fitting the characteristics of a BTS, suggesting a different subcellular localization. Nevertheless, both *E. longa* and *E. gracilis* have another homolog of this enzyme with an obvious BTS, so polyamines may be produced in the *E. longa* plastid after all (Fig. S4).

### *E. longa* plastid does not make fatty acids but maintains phospholipid and glycolipid synthesis

Eukaryotes synthesize fatty acids by a single multi-modular fatty acid synthase I (FASI) in the cytosol or by a multi-enzyme type II fatty acid synthesis complex in the plastid. *E. gracilis* possesses both systems (54), but *E. longa* encodes only a homolog of the cytosolic FASI enzyme (Fig. 2a; Table S5). Nevertheless, *E. longa* still maintains plastid-targeted versions of acyl carrier protein (ACP) and 4’-phosphopantetheinyl transferases (or holo-ACP synthase), which are crucial for the synthesis of an active form of ACP (55). This is apparently employed by the predicted plastid-targeted homologs of acyl-ACP synthetases (presumably activating fatty acids imported into the plastid) and enzymes required for the synthesis of phosphatidic acid (PA) and its subsequent conversion to phosphatidylglycerol (PG) (Fig. 2a; Table S5). Notably, *E. longa* also has a parallel, plastid-independent, route of phosphatidylglycerol synthesis (Table S6).

**Figure 2:**
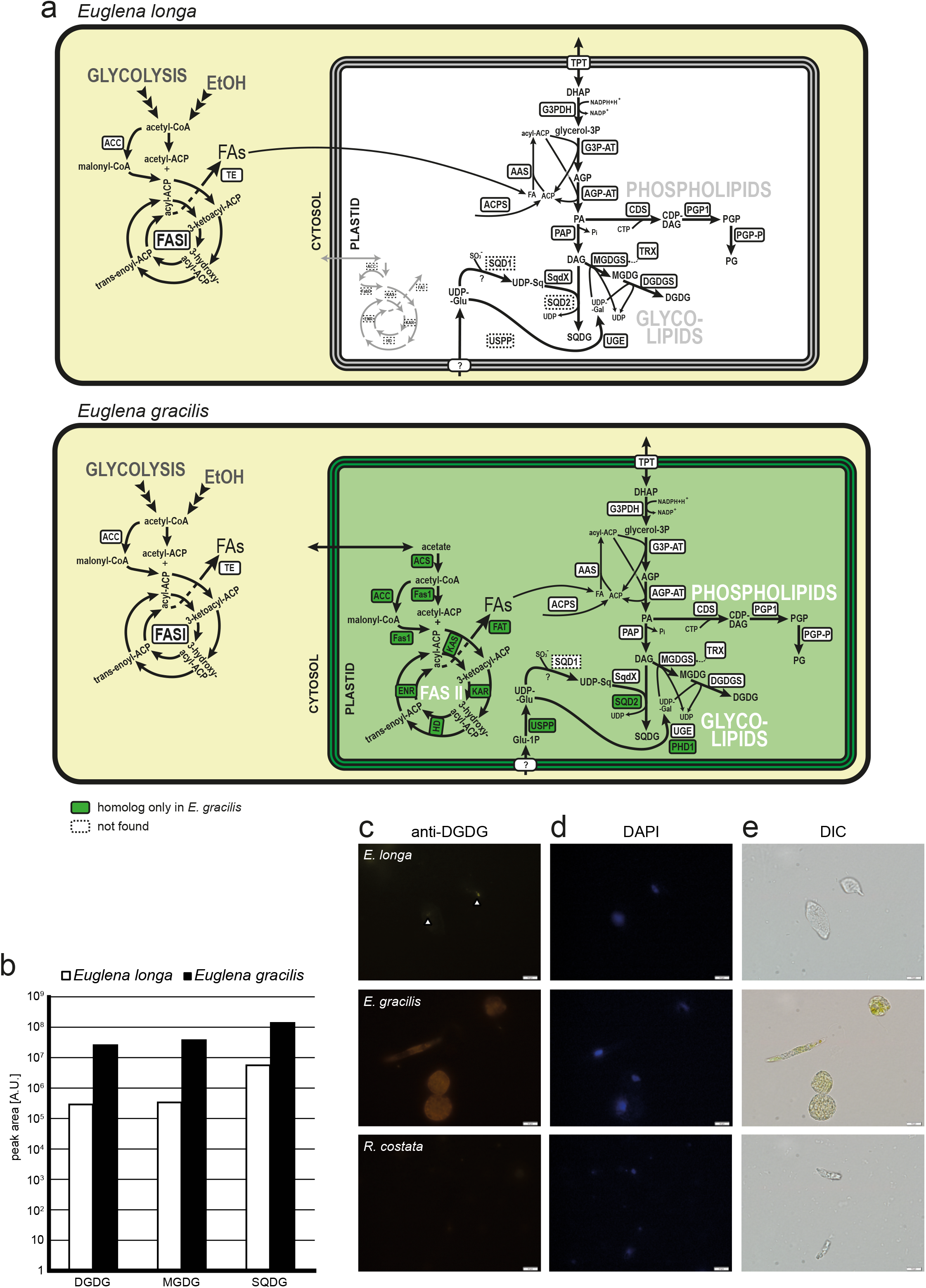
Fatty acid and lipid biosynthesis in *E. longa* and *E. gracilis.* **a**: Schematic comparison of the localization and evolutionary origin of enzymes. Abbreviations, fatty acid synthesis: ACC – acetyl-CoA carboxylase, ACS – acetyl-CoA synthetase, ENR – enoyl-CoA reductase, Fas1 – malonyl-CoA/acetyl-CoA: ACP transacylase, FASI – type I fatty acid synthase, FAT – fatty acyl-ACP thioesterase, HD – hydroxyacyl-ACP dehydratase, KAR – ketoacyl-ACP reductase, KAS – ketoacyl-ACP synthase, TE – fatty acid thioesterase, TRX – thioredoxin-regulated enzyme; glycolipid synthesis: AAS – acyl-ACP synthase, ACPS – holo-ACP synthase, AGP-AT – acylglycerophosphate acyltransferase, G3P-AT – glycerol-3-phosphate acyltransferase, G3PDH – glycerol-3-phosphate dehydrogenase, MGDG/DGDG – mono-/digalactosyl diacylglycerol, MGDGS/DGDGS – MGDG/DGDG synthase, PAP – phosphatidic acid phosphatase, SQD1 – UDP-sulfoquinovose synthase, SQD2/SQDX – sulfoquinovosyl diacylglycerol (SQDG) synthase, UGE/PHD1 – UDP-glucose epimerase, USPP – UDP-sugar pyrophosphorylase; phospholipid synthesis: CDS – CDP-diacylglycerol synthase, PGP1 – phosphatidylglycerophosphate synthase, PGP-P – phosphateidylglycerophosphate phosphatase. **b:** Semiquantitative comparison of glycolipids present in *E. longa* and autotrophically grown *E. gracilis.* Note the logarithmic scale of the quantification units (peak area). Peak area is an arbitrary unit expressing the intensity of the signal of a particular lipid species, recalculated according to their respective ionization promptitude. As each lipid species have different ionization promptitude, note that direct comparison can be done only within lipid class (for details, see Tomcala et al. 2017). **c-e:** Immunofluorescence micrographs using anti-DGDG antibody (C), DAPI (D) and differential interference contrast (E). Autotrophic *E. gracilis* represents a positive control, while the aplastidic euglenozoan *R. costata* was used as negative control.

No other reactions of phospholipid synthesis or decomposition beyond PG synthesis seem to operate in the *E. longa* plastid. However, enzymes for the synthesis of galactolipids monogalactosyldiacylglycerol (MGDG) and digalactosyldiacylglycerol (DGDG) were identified, all with predicted BTSs (Fig. 2a, Table S5), consistent with the plastidial localization of galactolipid synthesis in other eukaryotes (56). Moreover, both MGDG and DGDG could be detected in *E. longa* and *E. gracilis* by HPLC-MS/MS, although galactolipid levels were significantly lower in *E. longa* than in *E. gracilis* (Fig. 2b). The presence of DGDG was further confirmed by immunofluorescence using an anti-DGDG antibody, which showed DGDG to be present in small foci in the *E. longa* cells (Fig. 2c), presumably representing individual small plastids. In comparison, extensive staining was observed in *E. gracilis* cells consistent with plastids occupying a large portion of the cytoplasm, whereas no staining was observed in the plastid-lacking euglenid *Rhabdomonas costata.*

We additionally identified another typical plastid glycolipid, sulfoquinovosyldiacylglycerol (SQDG; 57) in both *Euglena* spp. (Fig. 2b). The enzyme directly responsible for SQDG synthesis is sulfoquinovosyltransferase (Fig. 2a), but interestingly, its standard eukaryotic version (SQD2) is present only in *E. gracilis*, whereas both species share another isoform phylogenetically affiliated to bacterial SqdX (Fig. 3). To our knowledge, this is the first encounter of SqdX in any eukaryote. The presence of SQD2 only in *E. gracilis* may relate to the specific needs of its photosynthetic plastid. Indeed, *E. gracilis* contains much more SQDG compared to *E. longa* (Fig. 2b), and the profile of esterified fatty acids differs between the two species *(E. longa* lacks SQDG forms with unsaturated longer chains; Table S7).

**Figure 3:**
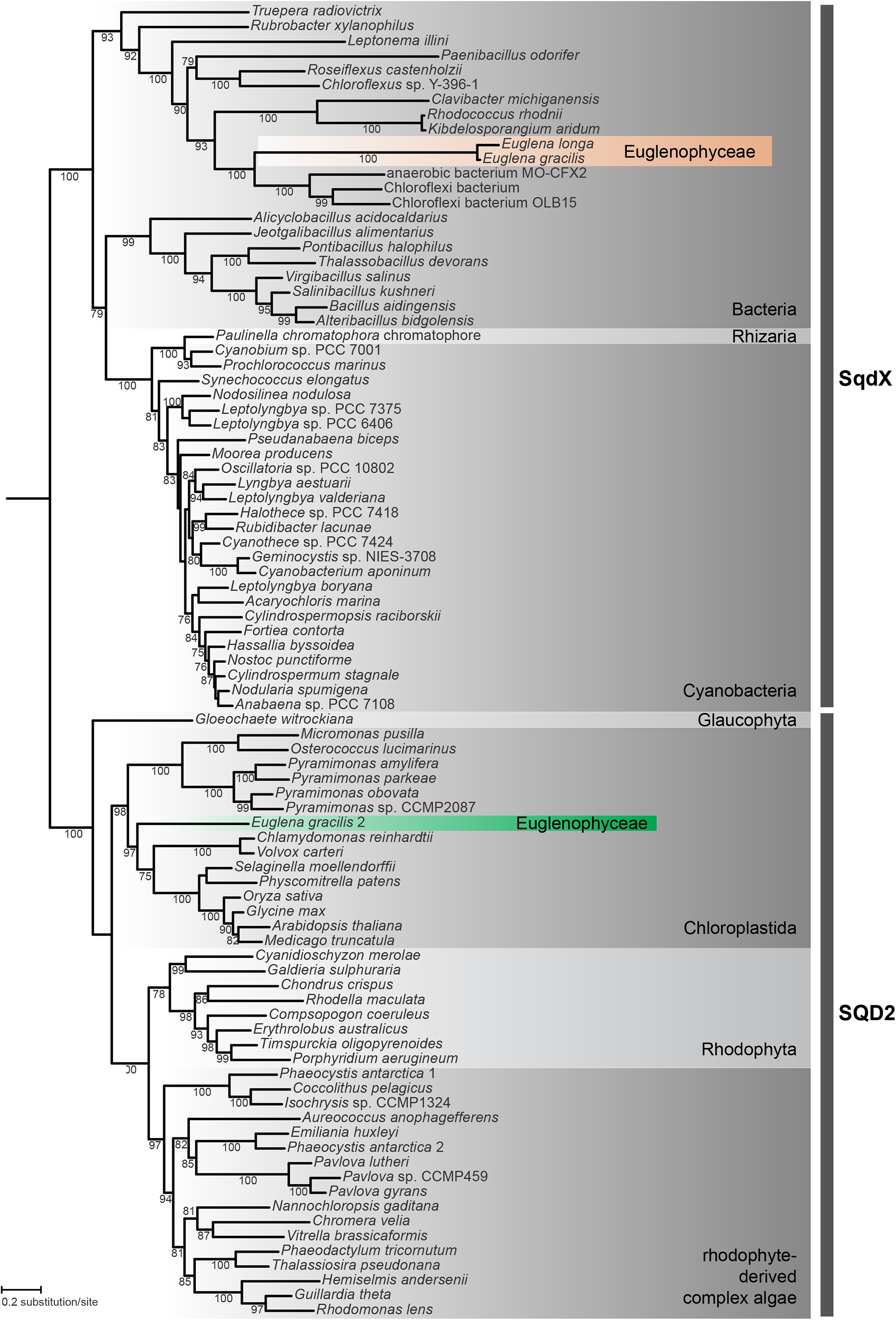
Euglenophytes have replaced the eukaryotic form of sulfoquinovosyltransferase (SQD2) with a bacterial version (SqdX). The maximum-likelihood tree was inferred with IQ-TREE using the LG+F+G4 substitution model and ultra-fast bootstrapping. The UFboot support values are indicated at branches when higher than 75%. Accession numbers of sequences included in the analysis are provided in Table S11.

The saccharide moieties of glycolipids in *E. longa* are probably also synthesized in its plastid (Fig. 2a). *E. longa* exhibits an ortholog of the E. *gracilis* UDP-glucose epimerase previously identified in the plastid proteome (Fig. S5, Table S5), explaining the source of UDP-galactose for galactolipid synthesis. This seems to be an original euglenozoan enzyme recruited into the plastid (Fig. S5), but, interestingly, *E. gracilis* encodes also a homolog of the unique plastidial UDP-glucose epimerase (PHD1) known from plants and various algae (58). The *E. gracilis* PHD1 possesses a predicted BTS (Table S5) and is thus also likely plastidial (albeit without proteomic support). This putative redundancy is not shared by *E. longa* (Fig. 2b) and may reflect a presumably much lower need for galactolipid synthesis. The origin of the SQDG precursor UDP-sulfoquinovose in *E. longa* remains obscure, because like *E. gracilis*, it lacks the conventional UDP-sulfoquinovose synthase SQD1/SqdB and probably employs an alternative, unrelated enzyme (38). UDP-glucose, i.e. the common precursor of both UDP-galactose and UDP-sulfoquinovose, may be produced directly in the plastid of *E. gracilis*, owing to the presence of an isoform of UDP-sugar pyrophosphorylase with a typical BTS (although absent among proteomically confirmed plastid proteins). *E. longa* lacks an ortholog of this protein as well as any other potentially plastidial enzyme of UDP-glucose synthesis (Table S5), suggesting import of this metabolite from the cytosol.

### *E. longa* plastid retains a linearized Calvin-Benson pathway

The expression of both subunits of RuBisCO in *E. longa* (32) raises the question whether the Calvin-Benson (CB) cycle (CBC) as a whole has been preserved in this organism. A putative *E. longa* plastid triose-phosphate isomerase was described previously (59), and we additionally identified homologs with putative BTSs for nearly all remaining CBC enzymes (Table S8). Phylogenetic analyses (supplementary dataset S1) showed specific relationships of the *E. longa* proteins to the previously characterized CBC enzymes from other euglenophytes (60). However, two key CBC enzymes are apparently missing from the *E. longa* transcriptome: phosphoglycerate kinase (ptPGK) and glyceraldehyde-phosphate dehydrogenase (ptGAPDH). Those homologs that are present are not orthologous to the plastid-targeted isoenzymes from other euglenophytes and all clearly lack a BTS (Table S8). Hence, these are presumably cytosolic enzymes involved in glycolysis/gluconeogenesis. The lack of ptPGK and ptGAPDH in *E. longa* implies that the product of the RuBisCO carboxylase activity, 3-phosphoglycerate (3PG), cannot be converted (via 1,3-bisphosphoglycerate; 1,3-BPG) to glyceraldehyde-3-phosphate (GA3P) in the plastid (Fig. 4a).

**Figure 4:**
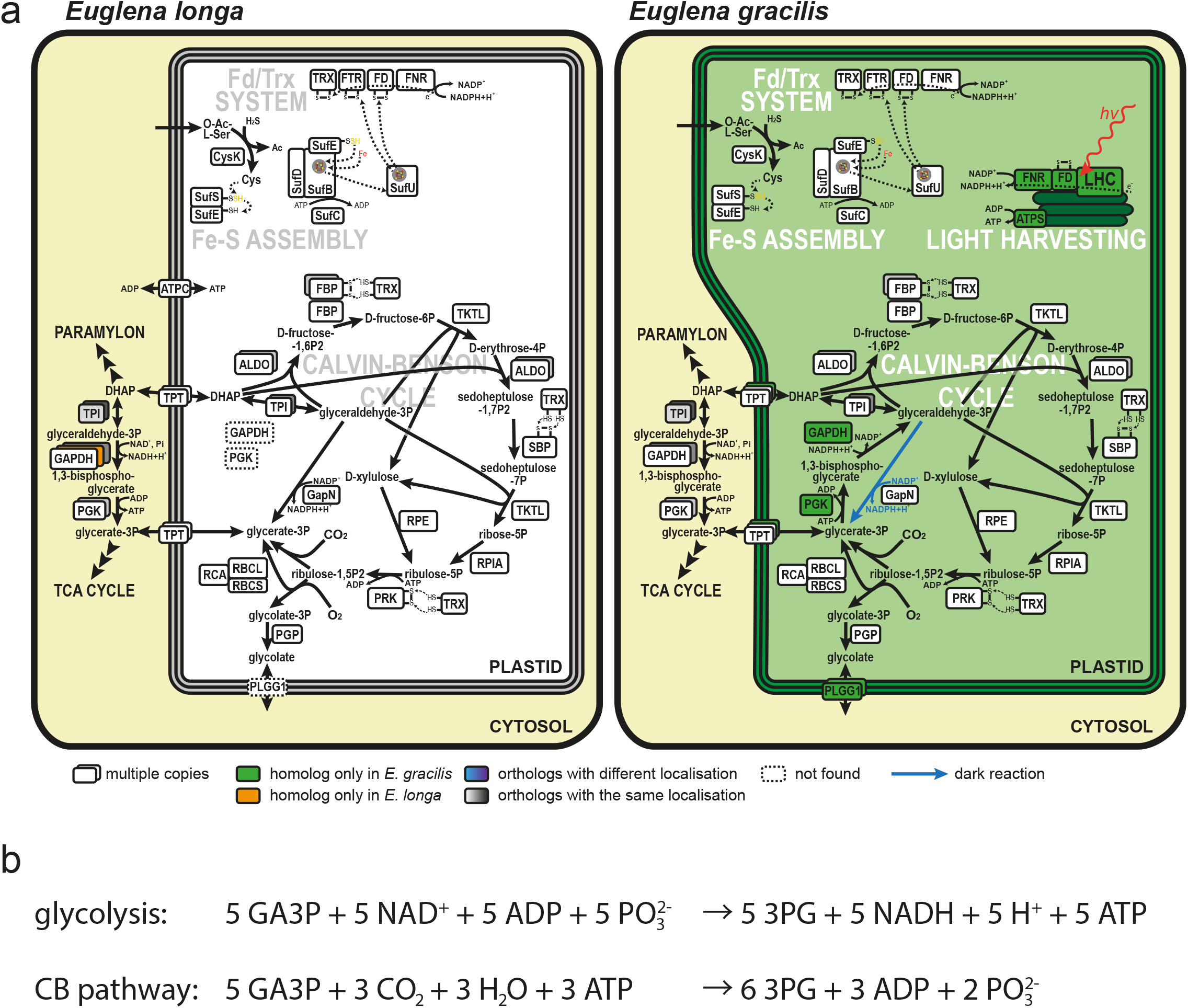
Carbon metabolism in the plastids of *E. longa* and *E. gracilis.* **a:** The Calvin-Benson cycle (CBC) resident to this organelle is central to the plastid carbon metabolism, regulated by the ferredoxin/thioredoxin (Fd/Trx) system. Reduction of disulfide bonds by the Fd/Trx system activates FBP and PRK. FTR and FD of the Fd/Trx system require for their function a post-translationally added Fe-S prosthetic group provided by the Fe-S assembly system. GapN apparently mediates shuttling of reducing equivalent (NADPH) through the exchange of DHAP/GA3P and 3PG, reflecting the cytosolic NADPH/NADP+ ratio and thus an overall metabolic state of the cell. In contrast, *E. gracilis* plastid is an energy-converting organelle, harvesting light into chemical energy bound as NADPH and ATP and subsequently using this bound energy to fix CO2 into organic carbohydrates via the CBC. Enzyme abbreviations are colour-coded according to their inferred evolutionary origin, see the graphical legend. **b:** Stoichiometric comparison of reactions converting glyceraldehyde 3-phosphate to 3-phosphoglycerate via glycolysis and the Calvin-Benson pathway. Abbreviations: 3PG – 3-phosphoglycerate, ALDO – aldolase, DHAP – dihydroxyacetone-phosphate, FBP – fructose-1,6-bisphosphatase, GA3P – glyceraldehyde-3- phosphate, GAPDH – glyceraldehyde-3-phosphate dehydrogenase, PGK – 3-phosphoglygerate kinase, PGP – phosphoglycolate phosphatase, PLGG1 – plastid glycolate/glycerate transporter, PRK – phosphoribulokinase, RBCL/RBCS – RuBisCO large/small subunit, RCA – RuBisCO activase, RPE – ribulose-5-phosphate epimerase; RPIA – ribulose-phosphate isomerase A, SBP – sedoheptulose-1,7-bisphosphatase, TKTL – transketolase, TPI – triose-phosphate isomerase, TPT – triose-phosphate translocator; Fd/Trx system: FD – ferredoxin; FNR – FD/NADP+ oxidoreductase, FTR – FD/TRX oxidoreductase, TRX – thioredoxin, ATPS – ATP synthase, ATPC – ADP/ATP translocase, LHC – light-harvesting complex.

Assuming that the reactions catalysed by fructose bisphosphatase, phosphoribulokinase, and RuBisCO are irreversible (61), the flux through this linearized CB pathway goes from GA3P to 3PG, with a net production of six molecules of 3PG from five molecules of GA3P due to fixation of three CO_2_ molecules catalysed by RuBisCO. Euglenophytes do not store starch in the plastid (62), and indeed, we did not find any glucose metabolism-related enzymes with a BTS in *E. longa.* Hence, GA3P cannot be produced by a glycolytic route in the *E. longa* plastid. The presence of the plastid-targeted glycerol-3-phosphate dehydrogenase (Table S5) in principle allows for generation of GA3P from glycerol-3-phosphate (via dihydroxyacetone phosphate; DHAP; Fig. 2), which could possibly come from glycerolipids turnover, but no plastidial phospholipid-degradation enzymes were found in *E. longa.* Hence, the primary function of glycerol-3-phosphate dehydrogenase perhaps is to provide glycerol-3 – phosphate for the plastid phospholipid and glycolipid synthesis (see above) and the *E. longa* plastid most likely imports GA3P or DHAP from the cytosol (Fig. 4a). This assumption is supported by the presence of several members of the plastid phosphate translocator (pPT) family (Fig. S6; 63), including one phylogenetically closest to a cryptophyte transporter with a preference for DHAP (64). Concerning the opposite end of the linear CB pathway, we did not identify any *E. longa* plastid-targeted enzyme that would metabolize 3PG further, suggesting that this intermediate is exported from the plastid into the cytosol, probably also by one of the pPT transporters (Fig. 4a).

RuBisCO is not only a carboxylase, but also exhibits an oxygenase activity catalysing the production of phosphoglycolate, which is then recycled by the photorespiration pathway; this is initiated by phosphoglycolate phosphatase, yielding glycolate (65). Indeed, E. *longa* contains an ortholog of the *E. gracilis* plastidial phosphoglycolate phosphatase (Table S8), but in contrast to *E. gracilis* no homolog of the glycolate transporter PLGG1 mediating glycolate export from the plastid (66) was found in *E. longa* (Table S8). Since it also lacks obvious candidates for plastid-targeted glycolate-metabolizing enzymes (glycolate oxidase, glyoxylate reductase, glycolaldehyde dehydrogenase, glyoxylate carboligase/tartronate-semialdehyde reductase), it is unclear how glycolate is removed from the *E. longa* plastid. Possibly the amount of glycolate produced is low and can be exported by an uncharacterized PLGG1-independent route that exists also in plant plastids (67) and is sufficient for glycolate recycling in the semi-parasitic plant *Cuscuta campestris* (68).

### *E. longa* plastid preserves the redox regulatory system of the CB pathway

Although the photosynthetic machinery is missing from *E. longa* (37), we found homologs (with clear plastidial localization) of the typical “photosynthetic” (PetF-related) ferredoxin (Fd) and ferredoxin-NADP+ reductase (FNR) (Table S9). These two proteins are primarily involved in passing electrons from activated photosystem I to NADP+. Euglenophyte FNR homologs belong to two different, yet related, clades (Fig. 5). One comprises the *E. longa* FNR plus its orthologs from photosynthetic euglenophytes, whereas the second one is restricted to the photosynthetic species. Two different FNR forms also exist in plants (Fig. 5), one functioning in photosynthesis (photosystem I-dependent production of NADPH) and the other “non-photosynthetic” one, allowing electron flow in the reverse direction, from NADPH to Fd (69). In analogy, we suggest that the two euglenophyte FNR forms functionally differ, one serving in photosynthesis and the other, present also in *E. longa*, mediating light-independent production of the reduced Fd. Multiple plastid anabolic enzymes depend on reduced Fd as an electron donor (4), but none of them seems to account for the presence of FNR and Fd in the *E. longa* plastid: glutamate synthase and nitrite reductase are missing, all identified lipid desaturases are predicted as mitochondrion- or ER-targeted (Table S5) and sulfite reductase, like the one previously identified in the plastid of *E. gracilis* (38), is NADPH-dependent (Table S5).

**Figure 5:**
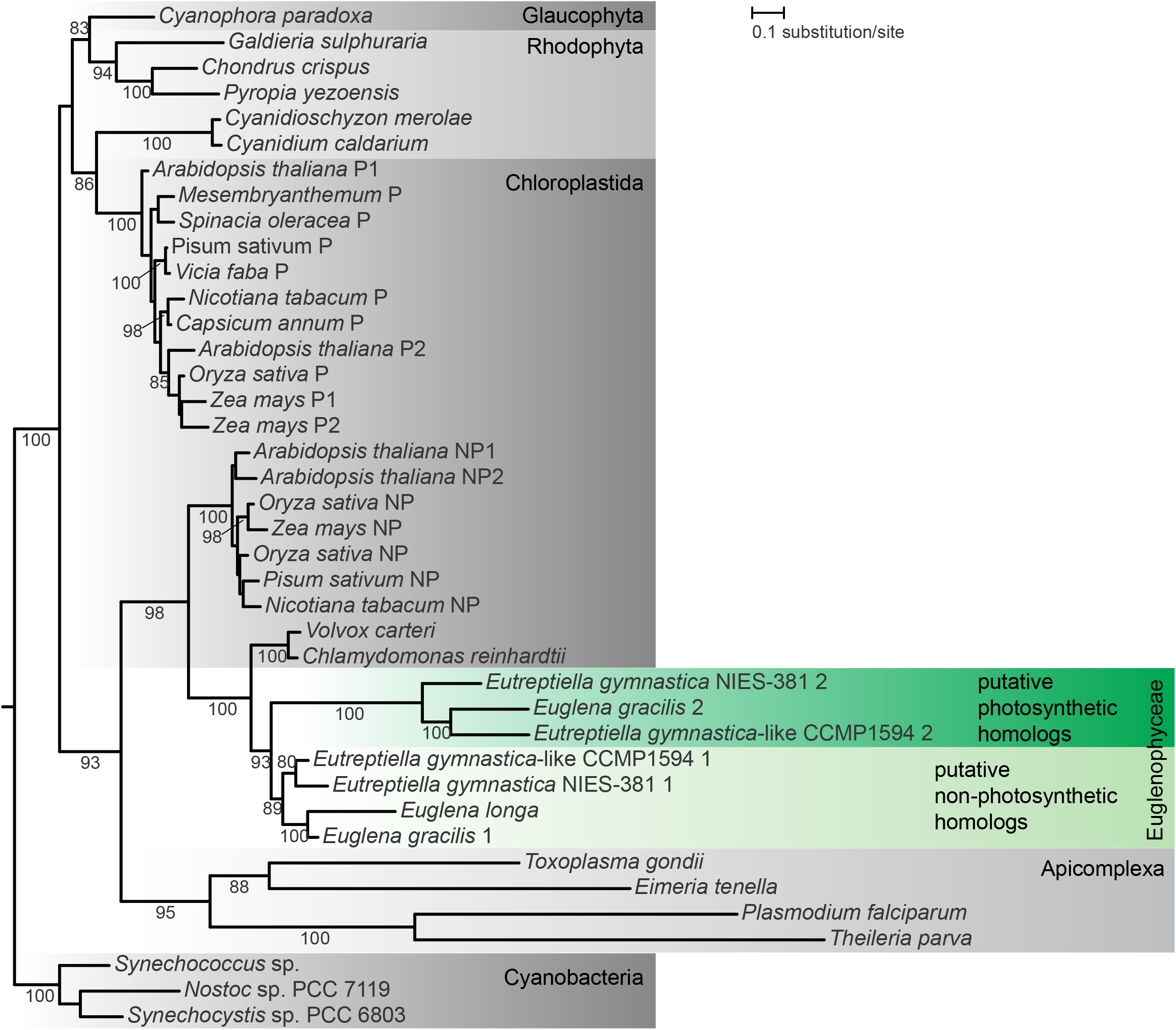
The inferred phylogeny of FNR. The maximum-likelihood tree was inferred with IQ-TREE using the LG+F+G4 substitution model and ultra-fast bootstrapping. The UFboot support values are indicated at branches when higher than 75%. Euglenophyte species are in bold, and their putative photosynthetic and non-photosynthetic homologs are depicted. The two forms of plant FNR are indicated: P, photosynthetic; NP, non-photosynthetic. Accession numbers of sequences included in the analysis are provided in Table S12.

Fd also provides electrons to ferredoxin:thioredoxin reductase (FTR) mediating reduction of the protein thioredoxin (Trx). The Fd/Trx system regulates several CBC enzymes in response to the stromal redox status, whereby an excess of NADPH leads to electrons being relayed from Fd *via* Trx to certain disulfide bonds in the target enzymes to activate them (Fig. 4a; 70). Notably, FTR and Trx homologs with an evident BTS are present in *E. longa* (Table S9), and specific motifs necessary for the function of the Fd/Trx system are conserved in the respective *E. longa* proteins (Fig. S7). In addition, three *E. longa* CBC enzymes, fructose bisphosphatase (two of the three isoforms present), sedoheptulose bisphosphatase, and phosphoribulokinase, exhibit the conserved Trx regulatory cysteine motifs, similar to their orthologs in *E. gracilis* (Fig. S7, Table S10). Thus, the *E. longa* CB pathway is likely to be sensitive to the redox status in the plastid, specifically to the concentration of NADPH (Fig. 4a).

## DISCUSSION

The analyses described above provide evidence for the cryptic *E. longa* plastid harbouring highly non-conventional combination of metabolic functions. Lacking the plastidial MEP pathway, *E. longa* joins the only recently discovered group of plastid-bearing eukaryotes with such a deficit, namely the colourless diatom *Nitzschia* sp. NIES-3581 (9) and various colourless chrysophytes (20, 21). An obvious explanation for this is that the cytosolic MVA pathway is sufficient to supply precursors for all cellular isoprenoids in these organisms. In contrast, the MEP pathway in apicomplexans and related alveolates (i.e. Myzozoa; 8), and in diverse non-photosynthetic chlorophytes (71), is essential, since the cytosolic MVA pathway was lost in these groups (72, 73). Strikingly, the *E. longa* plastid is still involved in isoprenoid metabolism, namely the synthesis of tocopherols and OH-PhQ. Like in *E. gracilis*, pathway leading to OH-PhQ cannot be presently reconstructed in full detail in either *Euglena* spp. (see also 38). Both euglenophytes studied lack homologs of the conventional enzymes of the middle part of the pathway (from o-succinylbenzoate to dihydroxynaphthoate) typically localized in the peroxisome (74). The respective enzyme activities were found associated with the plastid envelope in *E. gracilis* (75), suggesting an alternative solution that may hold for *E. longa,* too. The molecular identity of the putative PhQ hydroxylase (making OH-PhQ) is unknown, so its plastidial localization in *E. gracilis* or *E. longa* cannot be ascertained. Finally, a previously unknown step – reduction of the naphthoquinone ring – was demonstrated as a prerequisite for the reaction catalysed by MenG to proceed in plants and cyanobacteria (76). The respective reductase is well conserved among diverse cyanobacteria, algae and plants (74), but we did not identify close homologs in any of the euglenophyte transcriptome assemblies, suggesting that euglenophytes employ an unknown alternative enzyme.

*E. longa* seems to be the first eukaryote with a non-photosynthetic plastid documented to have retained the pathways for tocopherols and OH-PhQ synthesis. The presence of tocopherols in *E. longa* is not that surprising, as they are not restricted to photosynthetic tissues in plants and were detected also in non-photosynthetic *E. gracilis* mutants (42, 77). As potent lipophilic antioxidants, tocopherols might be employed by *E. longa* to protect its membrane lipids against reactive oxygen species generated by mitochondria and peroxisomes. The retention of OH-PhQ synthesis in *E. longa* is more puzzling, as the best-established role of (OH-)PhQ in plants and algae is its functioning as an electron carrier within the photosystem I (43, 78). PhQ was additionally proposed to serve as an electron acceptor required for proper function of photosystem II (79, 80). A homolog of the respective oxidoreductase (LTO1) is present in *E. gracilis* (Table S2), but not in the transcriptomic data from *E. longa.* Interestingly, in plants PhQ was also detected in the plasma membrane and proposed to be involved in photosynthesis-unrelated redox processes (81–83). However, the MenA and MenG enzymes in *E. longa* carry a typical BTS, so we suggest that OH-PhQ in *E. longa* is involved in a hitherto uncharacterized, photosynthesis-unrelated plastid-resident process.

The absence of the type II fatty acid synthesis in the *E. longa* plastid is noteworthy, yet not unprecedented, since it has been also reported for the non-photosynthetic plastids of certain myzozoans (8) and a chrysophyte (20). Still, the *E. longa* plastid plays an active role in the lipid metabolism, having retained biosynthesis of several glycerolipid types, including galactolipids and SQDG. These were previously documented in several non-photosynthetic algae, e.g. colourless diatoms (84, 85). On the other hand, the apicoplast (86, 87), and most likely also the relic plastid of *Helicosporidium* (based on our analysis of the respective genome data reported by 17), lack galactolipid and SQDG synthesis completely. The reason for the differential retention of these lipids in different colourless plastids remains to be investigated further.

The truly striking feature of the *E. longa* plastid is the retention of nearly all CBC enzymes (assembling a putative linear CB pathway) and the mechanism of their redox regulation. In fact, the presence of CBC enzymes have been reported from a set of unrelated colourless algae and plants. Some of them, e.g. the dinoflagellate *Crypthecodinium cohnii,* dictyochophytes *Pteridomonas danica* and *Ciliophrys infusionum,* the cryptophyte *Cryptomonas paramecium,* and some parasitic or mycoheterotrophic land plants, are known to encode RuBisCO (7, 15, 88–90), but the actual complement of other CBC enzymes in these species is unknown. In contrast, transcriptomic or genomic analyses of other colourless plastid-bearing taxa, such as the dinoflagellate *Pfiesteria piscicida*, the chlorophyte *Helicosporidium* sp. ATCC50920, the diatom *Nitzschia* sp. NIES-3581, and the non-photosynthetic chrysophytes, revealed the presence of a subset of CBC enzymes, including ptPGK and ptGAPDH, but not of RuBisCO (9, 17, 21, 91). Hence, the constellation of the CBC enzymes present in the *E. longa* plastid is unique.

The CBC enzymes retained in various non-photosynthetic eukaryotes obviously do not serve to sustain autotrophic growth due to lack of photosynthetic production of ATP and NADPH. The incomplete CBC in *Nitzschia* was proposed to provide erythrose-4-P for the synthesis of aromatic amino acid *via* the shikimate pathway (9). The data provided for the *Helicosporidium* plastid (17) offer the same explanation of the retention of several CBC enzyme. However, such rationalization cannot hold for *E. longa,* since aromatic amino acid biosynthesis in this species apparently localizes to the cytosol (Table S3) and thus has an access to erythrose-4-P produced by the pentose phosphate pathway. In addition, *E. longa* differs from both *Nitzschia* and *Helicosporidium* by the retention of RuBisCO. A photosynthesis- and CBC-independent role of RuBisCO was described in oil formation in developing seeds of *Brassica napus,* where refixation of CO2 released during carbohydrate-to-fatty acid conversion increases carbon use efficiency (92). The absence of fatty acid synthesis in the *E. longa* plastid makes a similar function of RuBisCO unlikely in this organism.

The identification of the Fd/Trx system in the *E. longa* plastid, despite the absence of photosynthesis, may thus be a key for understanding the physiological role of the linear CB pathway in *E. longa.* Another hint is provided by the discovery of a unique (non-phosphorylating) form of GAPDH, referred to as GapN, in the *E. gracilis* plastid (38). This enzyme uses NADP+ to directly oxidize GA3P to 3PG without ATP generation (93). In plants, GapN is cytosolic and involved in shuttling of reducing equivalents from the plastid by the exchange of GA3P and 3PG between the two compartments (94). *E. longa* possesses a protein orthologous to the *E. gracilis* GapN with predicted BTS (Table S8), suggesting its plastidial localization. It thus appears that in *Euglena* spp., GapN mediates shuttling of reducing equivalents in the opposite direction than in plants, i.e. from the cytosol to the plastid (Fig. 4a). In case of *E. longa* this may be the main (if not the only) mechanism of providing NADPH for the use in the plastid, whereas *E. gracilis* would utilize it when photosynthetic NADPH production is shut down. At the same time, the shuttle provides a mechanism of linking the level of NADPH in the plastid with the cytosolic concentration of GA3P.

Taken together, we propose that in *E. longa* (and, at specific circumstances, possibly also in *E. gracilis),* the plastidial NADPH/NADP+ ratio is directly influenced by the redox status of the cell, i.e. that it rises in an excess of reducing power that slows down the glycolytic oxidation of GA3P in the cytosol. This stimulates the linear CB pathway *via* the Fd/Trx system, effectively decreasing the level of GA3 by converting it to 3PG without further increasing the reducing power in the cell. This conclusion is apparent from considering the overall stoichiometries of the two alternative pathways from GA3 to 3PG (Fig. 4b). The key difference is that the CB pathway does not produce NADH that needs to be reoxidized to keep the glycolytic pathway running, since the fixed CO2 effectively serves as an electron acceptor. Hence, turning the CB bypass on may help the cell to keep the redox balance when reoxidation of NADH is not efficient, e.g. at hypoxic (or anoxic) conditions (although this happens at the expense of ATP). Indeed, euglenophytes in their natural settings are probably often exposed to the shortage of oxygen, and anaerobiosis in *E. gracilis* has been studied to some extent (54, 95). The anaerobic heterotrophic metabolism of *E. gracilis* relies on fermentative degradation of paramylon leading to production of wax esters (96). It is likely that *E. longa* exhibits a similar metabolic adaptation to low oxygen levels as *E. gracilis.* However, details of the euglenophyte anaerobic metabolism need to be investigated further, and we propose that the plastid may be involved in it as a “redox valve”.

Compared to the range of forms mitochondria may exhibit in diverse eukaryotes (97), plastids seem to be much more uniform. However, this is partly a reflection of our ignorance about plastid biology in most algal groups, and recent studies of various independently evolved colourless plastids document a surprising degree of diversity in terms of their metabolic capacity. Our analyses of the *E. longa* plastid stretch the breadth of variation among non-photosynthetic plastids even further. The combination of pathways present (tocopherol and phylloquinone synthesis, glycolipid synthesis and a linearized CB pathway including RuBisCO), absent (fatty acid, amino acid, and isoprenoid precursor synthesis), and truncated (tetrapyrrole synthesis; Füssy, Záhonová, Oborník & Eliáš, unpublished) makes the *E. longa* plastid unlike any of the previously investigated non-photosynthetic plastids, including the apicoplast. However, further work, combining additional *in silico* analyses (aimed, e.g., at potential plastid membrane transporters mediating metabolite exchange with the cytosol) with biochemical and cytological investigations is needed to achieve a more precise idea about the protein composition of the *E. longa* plastid and a better understanding of its physiological roles.

## MATERIALS AND METHODS

### Identification and annotation of plastid-targeted proteins

The analyses utilized the *E. longa* transcriptome assembly reported previously, with candidates for plastid-targeted proteins identified as described in (37), including careful manual curation of the sequences and, if needed, revision of the 5’-ends of the transcripts by targeted searches of unassembled sequencing reads. Protein models with a putative BTS were automatically annotated using InterProScan 5.21 (98). Potential plastid enzymes (references from the KEGG PATHWAY Database, https://www.genome.jp/kegg/pathway.html) or sequences identified by literature searches, and plastid proteins identified by (38) were searched using BLAST v.2.2.30 (against the conceptually translated proteome, the transcriptome assembly and RNA-seq reads). HMMER 3.0 (99) was used when BLAST did not yield expected candidate homologs. For comparative purposes we used the same approach to identify plastid-targeted proteins encoded by the transcriptome assemblies from *E. gracilis* reported by (96) (accession GDJR00000000.1) and (100) (accession GEFR00000000.1).

To identify orthologs of the proteins from *E. gracilis* plastid proteome (38) in *E. longa,* these were used as queries in reciprocal BLAST searches. Briefly, *E. gracilis* proteins identified in its plastid proteome were used as queries in tBLASTn searches in *E. longa* transcriptome with E-value cut-off 0.1. Each respective best BLAST hit from *E. longa* was then used as a query to search the whole *E. gracilis* transcriptomic database and was classified as an ortholog if it retrieved the original *E. gracilis* query as a first hit. Results are summarized in Table S1.

For MenA cDNA resequencing, mRNA was extracted using the TRI Reagent and Dynabeads mRNA Purification kit (both from Thermo Fisher Scientific, Waltham, USA). Reverse-transcription was performed with random hexamers and StrataScript III Reverse Transcriptase (Thermo Fisher Scientific). The target was amplified using forward 5’-GGTGCTGTTCTGCTCTCACT-3’ and reverse 5’-CAGTGGGGATCAGAGATGCG-3’ primers, and Q5 High-Fidelity DNA polymerase in a standard buffer (New England Biolabs, Ipswich, USA). Amplicons were purified on MinElute PCR Purification columns (Qiagen, Hilden, Germany) and sequenced at the GATC sequencing facility (Konstanz, Germany). The MenA cDNA sequence is deposited in GenBank (MK484704).

### Phylogenetic analyses

Homologs of target proteins were identified by BLAST v.2.2.30 searches in the non-redundant protein sequence database at NCBI (www.ncbi.nlm.nih.gov) and among protein models of selected organisms from JGI (jgi.doe.gov) and MMETSP (marinemicroeukaryotes.org; 101). Datasets were processed using an in-house script (https://github.com/morpholino/Phylohandler) as follows. Sequences were aligned using the MAFFT v7.407 tool with L-INS-I setting (102) and poorly aligned positions were eliminated using trimAl v1.4.rev22 with “-automated1” trimming (103). For presentation purposes, alignments were processed using CHROMA (104). Maximum likelihood trees were inferred using the LG+F+G4 model of IQ-TREE v1.6.9 (105), employing the strategy of rapid bootstrapping followed by a “thorough” likelihood search with 1,000 bootstrap replicates. The list of species, and the number of sequences and amino acid positions are present in Tables S11-S22 for each phylogenetic tree.

### Culture conditions

*Euglena gracilis* strain *Z* (“autotrophic” conditions) was cultivated statically under constant illumination at 26 °C in Cramer-Myers medium with ethanol (0.8% v/v) as a carbon source (106). *E. longa* strain CCAP 1204-17a (a gift from Wolfgang Hachtel, Bonn, Germany) and heterotrophic E. *gracilis* strain *Z* were cultivated as above, but without illumination. *Rhabdomonas costata* strain PANT2 (a gift from Vladimír Hampl, Charles University, Prague, Czech Republic) was isolated from a freshwater body in Pantanal (Brazil) and grown with an uncharacterised mixture of bacteria in Sonneborn’s *Paramecium* medium, pH 7.4 (107) at room temperature.

### Mass spectrometry of structural lipids and terpenoids

Lipid extracts from *E. longa* and autotrophically grown *E. gracilis* cellular pellets (four biological samples of different culture ages) were obtained with procedures described in (108). Briefly, approximately 10 mg (wet weight) of both harvested cultures were homogenized by using a TissueLyser LT mill (Qiagen) and extraction was performed by chloroform and methanol solution (ratio – 2:1) following the previously described method (109). Aliquots from each sample were subjected to HPLC MS system powered by a linear ion trap LTQ-XL mass spectrometer (Thermo Fisher Scientific). The settings of the system followed the methodology published earlier (108). Data were acquired and processed using Xcalibur software version 2.1 (Thermo Fisher Scientific). Particular compounds were determined based on earlier publication (108). Terpenoids were extracted from an autotrophic and heterotrophic culture of *E. gracilis,* and a culture of *E. longa* of the same age in three replicates. The same extraction protocol as for lipid analysis was used. Sample aliquots were injected into the high-resolution mass spectrometry system powered by Orbitrap Q-Exactive Plus with Dionex Ultimate 3000 XRS pump and Dionex Ultimate 3000 XRS Open autosampler (both from Thermo Fisher Scientific) and followed the settings described in (108). Data were acquired and processed using Xcalibur software version 2.1. Identification of OH-PhQ was done by considering the m/z value, fragmentation pattern, and high-resolution data. Tocopherols (α, β/γ, and δ) were determined by the same characteristics as OH-PhQ and results were then compared with commercially purchased standards (Sigma-Aldrich, St. Louis, USA).

### Immunofluorescence assay

Immunofluorescence was performed as described previously (110). Briefly, cells were fixed in 4% paraformaldehyde for 30 minutes, permeabilized for 10 minutes on ice with 0.1% Igepal CA-630 (Sigma-Aldrich) in PHEM buffer pH 6.9 (60 mM PIPES, 25 mM HEPES, 10 mM EGTA, 2 mM MgCl_2_), and background was masked with 3% BSA in PHEM buffer. DGDG was detected using a polyclonal rabbit anti-DGDG antibody (1:25), a kind gift from Cyrille Y. Botté (University of Grenoble I, Grenoble, France), followed by incubation with a secondary Cy3-labeled polyclonal goat anti-rabbit antibody (AP132C, 1:800, Merck Millipore, Burlington, USA). Cells were mounted on slides using Fluoroshield™ with DAPI mounting medium (Sigma-Aldrich) and observed with an Olympus BX53 microscope (Olympus, Tokyo, Japan).

## Supporting information

Supplementary Data - Newick trees

## ACKNOWLEDGEMENTS

We thank Vladimír Hampl for the culture of *Rhabdomonas costata* and Cyrille Y. Botté for the anti-DGDG antibody. We acknowledge the infrastructure grant “Přístroje IET” (CZ. 1.05/2.1.00/19.0388). Computational resources were supplied by the Ministry of Education, Youth and Sports of the Czech Republic under the Projects CESNET (Project No. LM2015042) and CERIT-Scientific Cloud (Project No. LM2015085) provided within the program Projects of Large Research, Development and Innovations Infrastructures. We thank Laboratory of Analytical Biochemistry and Metabolomics (Biology Centre ASCR) for an access to LC-MS instruments. This study was supported by the Czech Science Foundation grants 17-21409S (to ME) and 18-13458S (to MO), ERD Funds (the project CePaViP; CZ.02.1.01/0.0/0.0/16_019/0000759) and the Scientific Grant Agency of the Slovak Ministry of Education (grant VEGA 1/0535/17 to JK).

## Notes

### Competing Interest Statement

The authors have declared no competing interest.

