## Supplementary Figures S1-S7 for "The cryptic plastid of *Euglena longa* defines a new type of non-photosynthetic plastid organelles"

**Figure S1: Inferred phylogeny of phytol kinase VTE 5 and phytyl-phosphate kinase VTE6.** The maximum-likelihood tree was inferred with IQ-TREE using the LG+F+G4 substitution model and ultra-fast bootstrapping. The UFboot support values are indicated at branches when higher than 75%. Colour of the Euglenophyceae clade reflects possible origin in Chloroplastida (green) or Rhodophyta-derived complex algae (ochre).

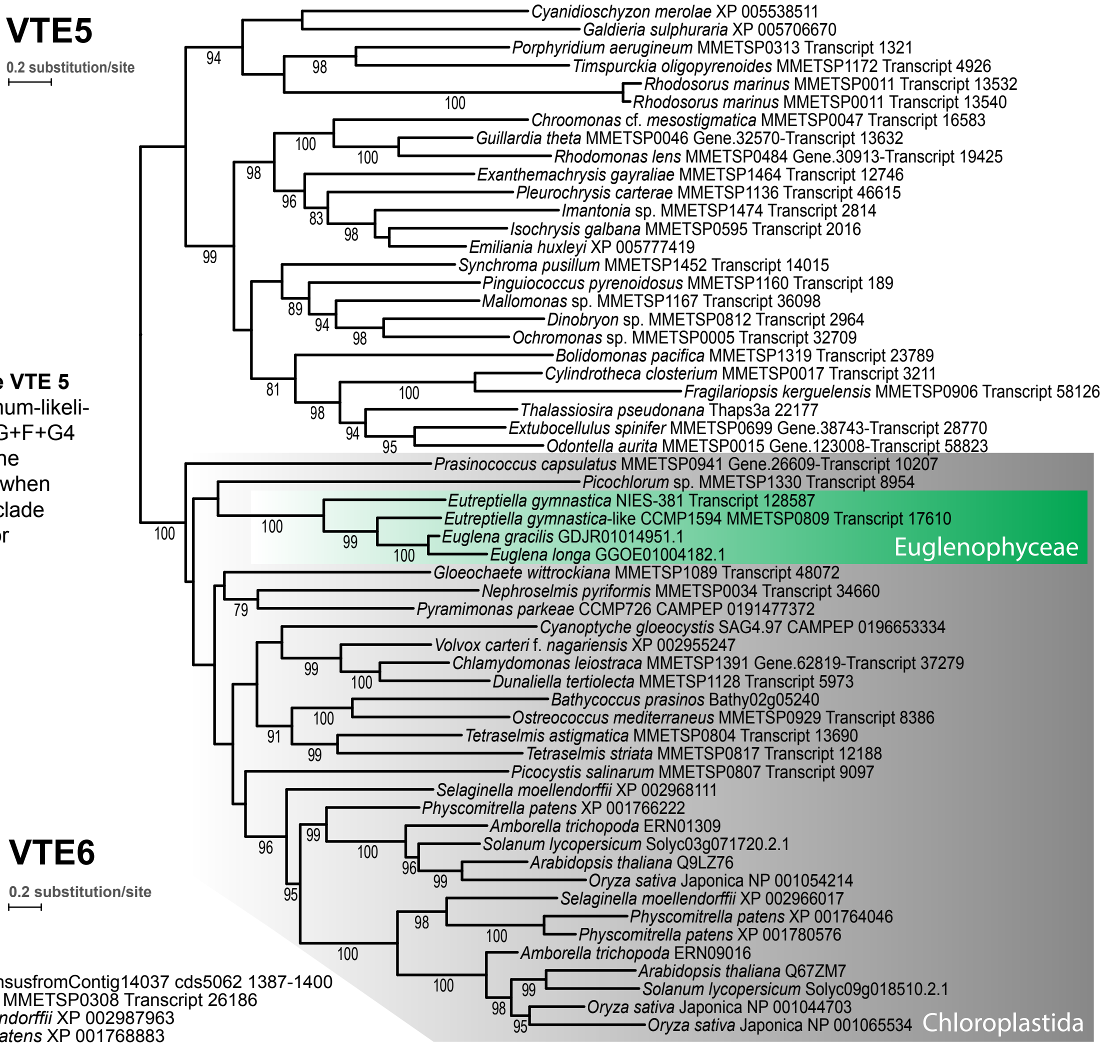


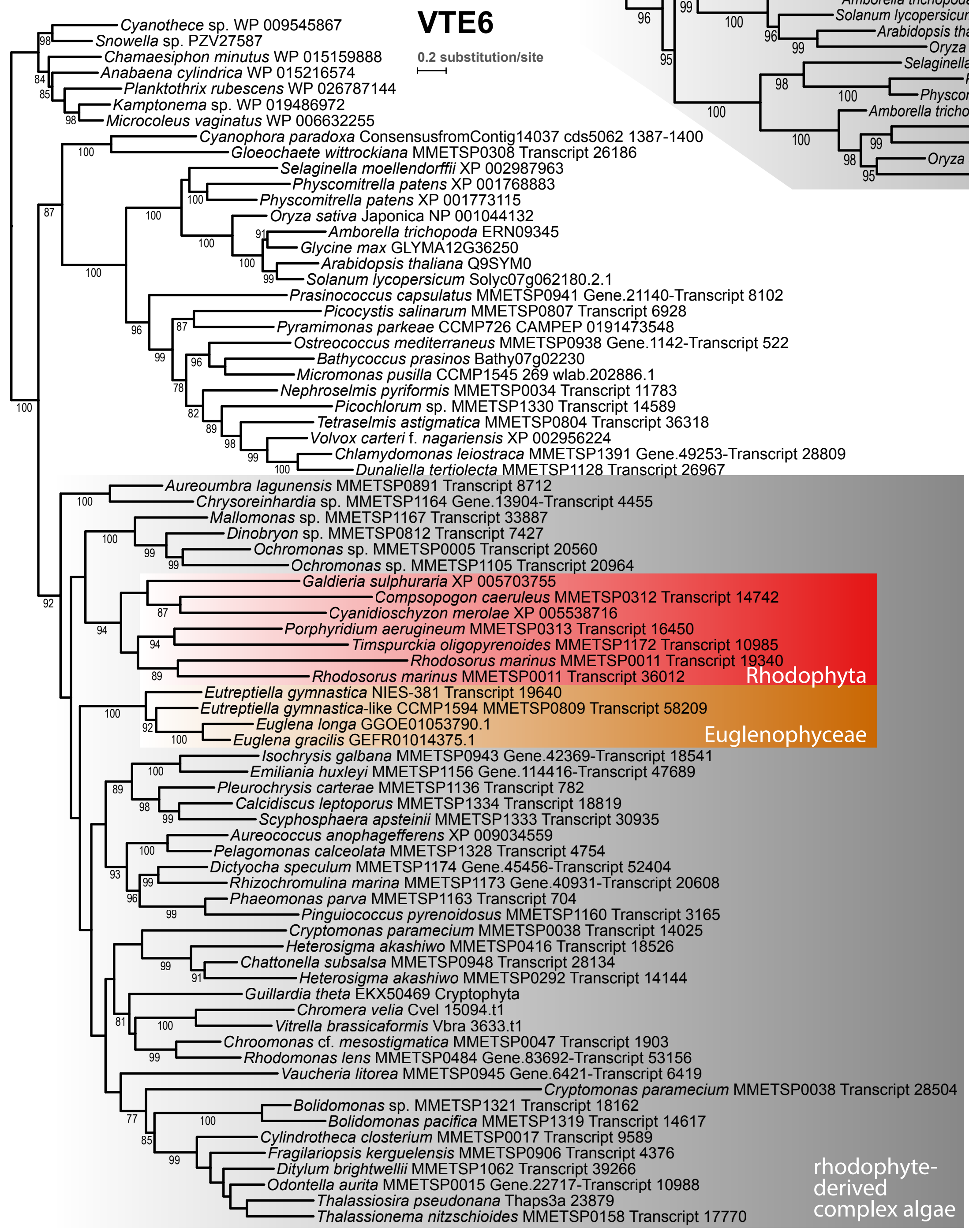

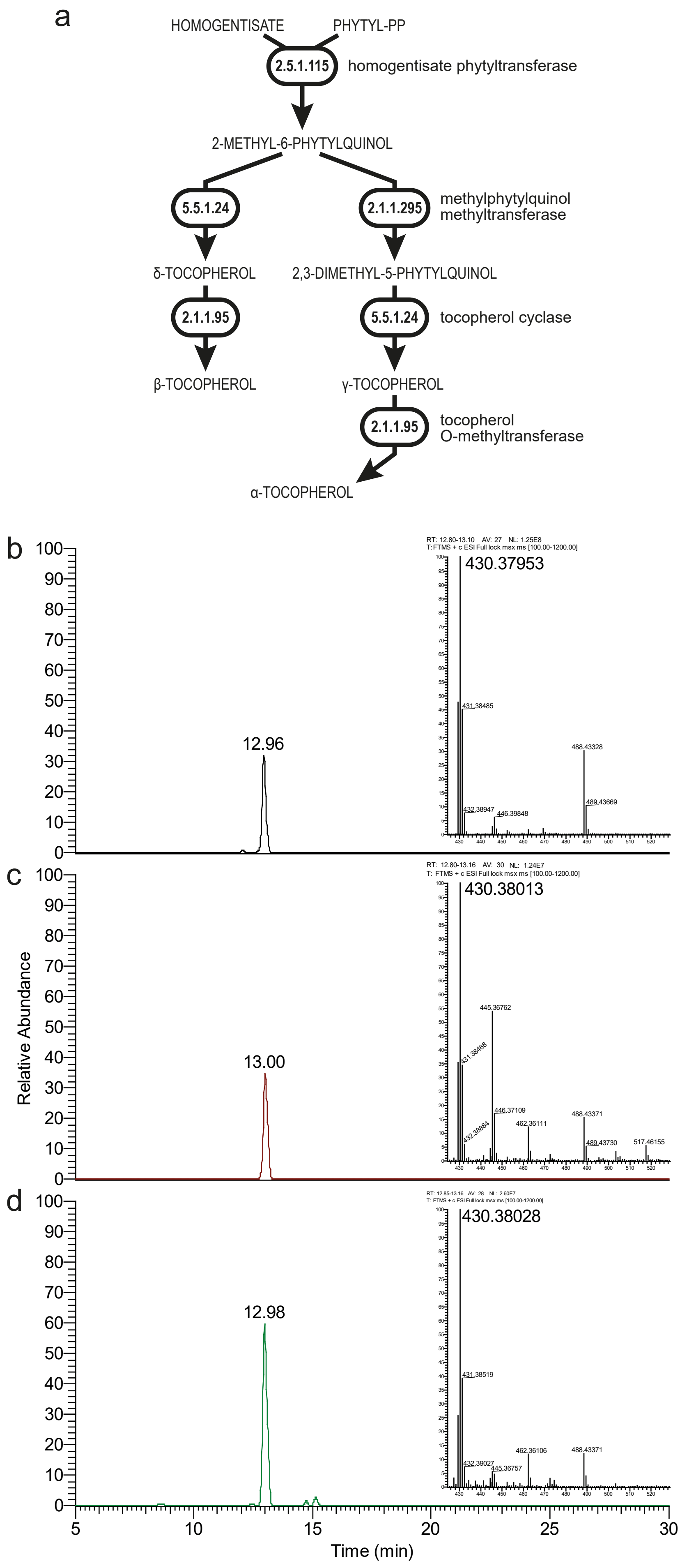

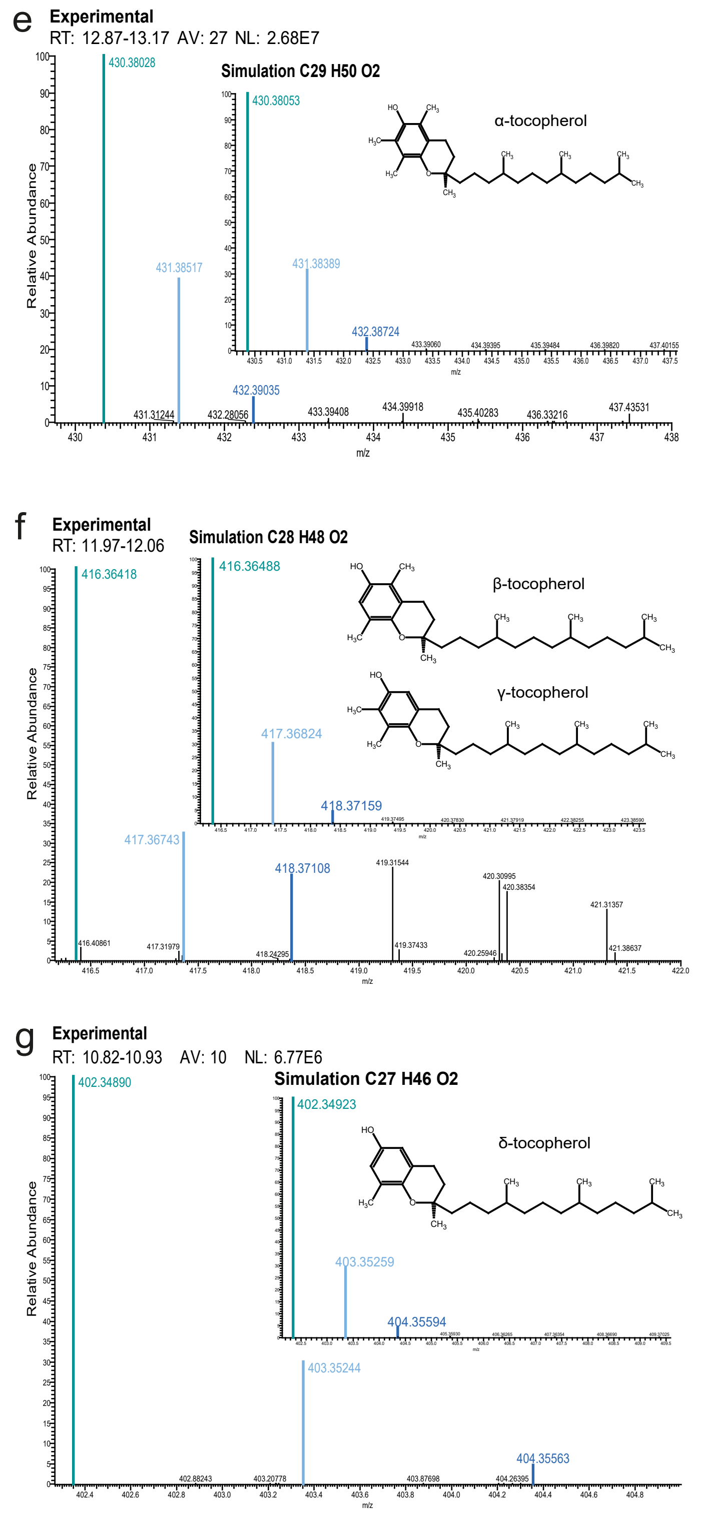


h

| | | α-tocopherol | β-/γ-tocopherol | δ-tocopherol |
|---|---|---|---|---|
| simulation | [M]+[Da] | 430.38052 | 416.36488 | 402.34923 |
| standards | | 430.37947 | 416.36443 | 402.34897 |
| *E. gracilis* heterotrophic | | 430.38343 | 416.36440 | 402.35766 |
| *E. longa* | | 430.38053 | 416.36418 | 402.34890 |
| standards | Δ [M]+[Da] | -0.00105 | -0.00045 | -0.00026 |
| *E. gracilis* heterotrophic | | 0.00396 | -0.00048 | 0.00843 |
| *E. longa* | | -0.00290 | -0.00070 | -0.00033 |

**Figure S2: Experimental detection of tocopherols in *E. longa* and *E. gracilis*. a:** Overview of tocopherol biosynthesis (enzymes indicated by their EC numbers). **b-d:** Extracted chromatograms of exact mass of α-tocopherol (m/z 430.3805; inset) and recorded spectra from a particular peak in an α-tocopherol standard (b), raw lipid extracts of heterotrophic *E. gracilis* (c) and *E. longa* (d). **e-g:** Comparison of simulation of α- (e), β- and γ- (f), and δ-tocopherol (g) chemical formula spectrum and experimentally gained data from the *E. longa* sample. The chemical structure of the particular tocopherol is shown as inset. **h:** Comparison of tocopherols' monoisotopic mass simulated and obtained high-resolution data in examined euglenophyte samples.

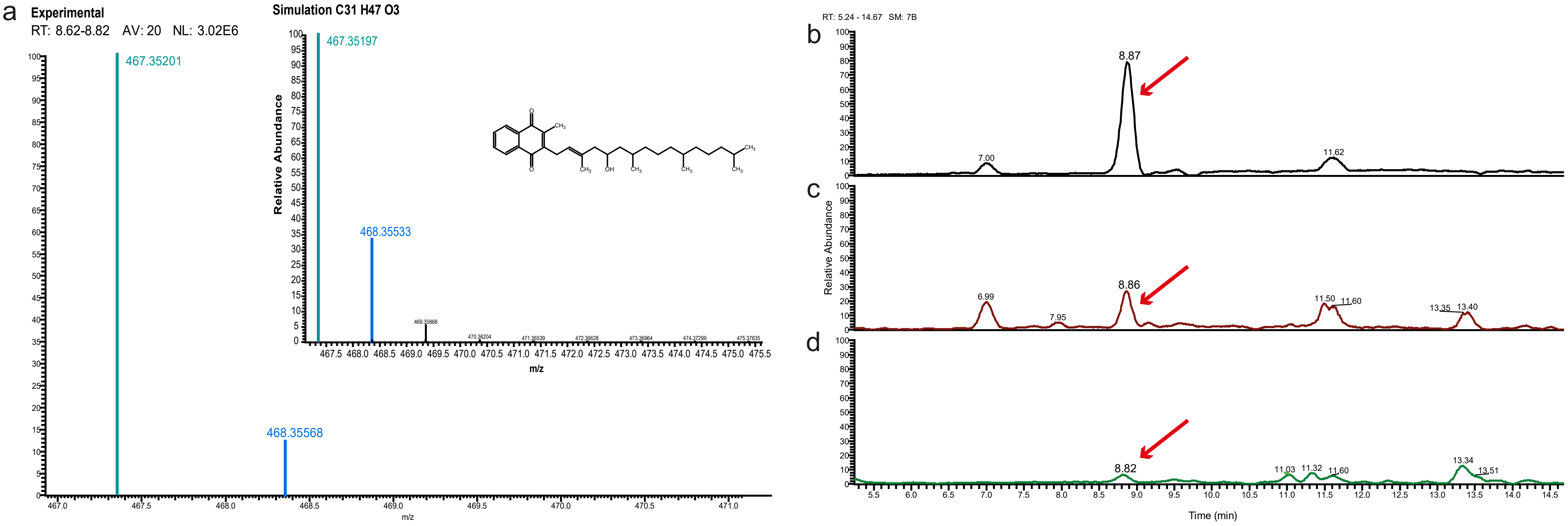


e

| 5 - hydroxyphylloquinone | $[M+H]^+$ [Da] | Δ $[M+H]^+$[Da] |
|---|---|---|
| simulation | 467.35197 | |
| *E. gracilis* autotrophic | 467.35191 | 0.00006 |
| *E. gracilis* heterotrophic | 467.35196 | 0.00001 |
| *E. longa* | 467.35202 | -0.00005 |

**Figure S3. Experimental confirmation of 5-hydroxyphylloquinone (OH-PhQ) in *E. longa* and *E. gracilis*. a:** Comparison of simulation of OH-PhQ chemical formula spectrum and experimentally gained data from autotrophic *E. gracilis* sample. The chemical structure of OH-PhQ is shown as inset. **b-d:** Extracted chromatograms of exact mass of protonated OH-PhQ (m/z 467.35) of raw lipid extracts. The red arrow points to a peak of OH-PhQ determined by high-resolution and fragmentation pattern in autotrophic *E. gracilis* (b), heterotrophic *E. gracilis* (c), and *E. longa* (d). **e:** Comparison of OH-PhQ monoisotopic mass simulated and obtained high-resolution data in examined euglenophyte samples.

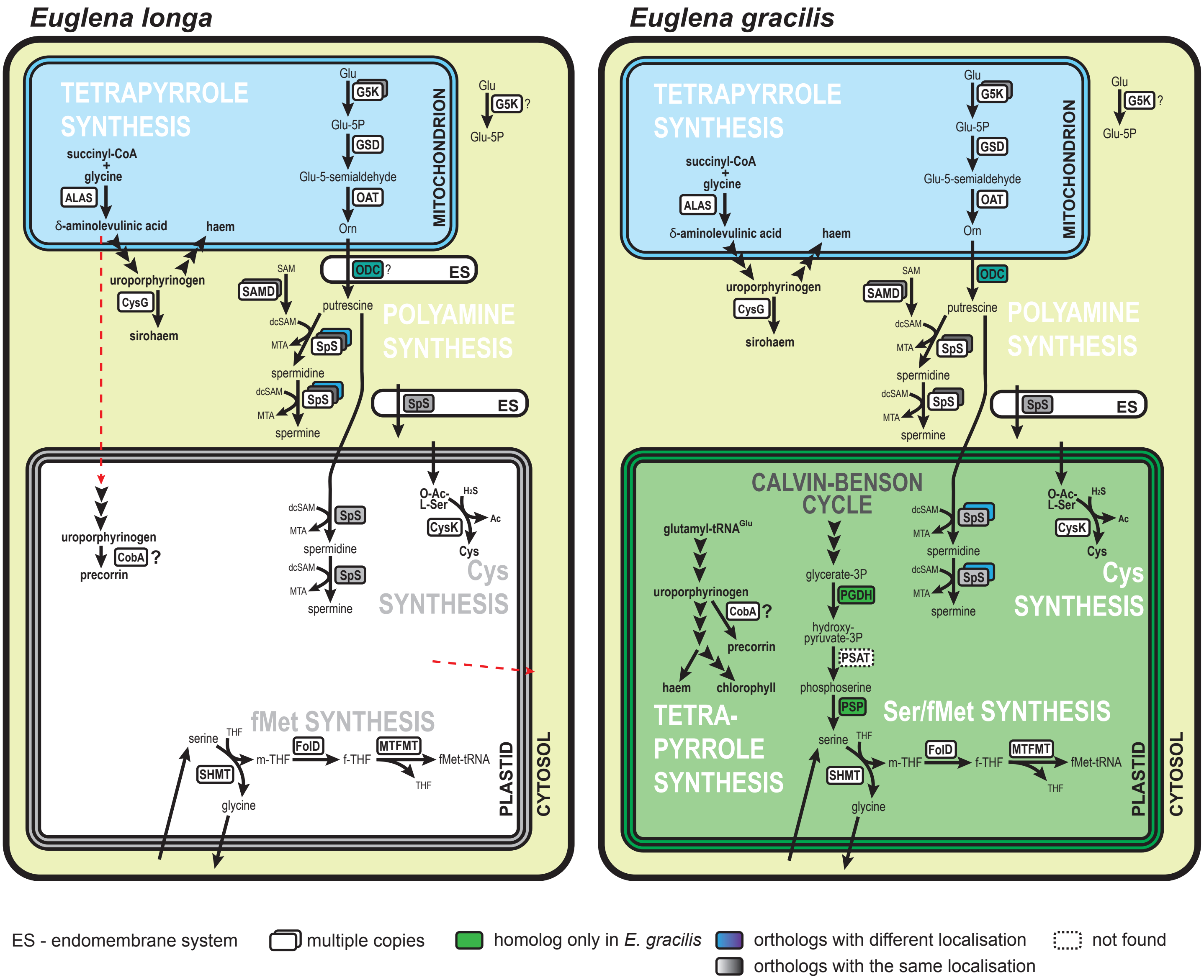


**Figure S4: Plastid-linked nitrogen metabolism in *E. longa* and *E. gracilis*.** Schematic comparison of the localization of enzymes of plastid tetrapyrrole, serine, formylmethionine, cysteine and polyamine synthesis. For simplicity, triple arrowheads represent multiple enzymatic/transport steps in a depicted pathway. Red arrow indicates a proposed intermediate transport to support plastid precorrin synthesis in *E. longa* (details elaborated elsewhere). Abbreviations, ALAD – delta-aminolevulinic acid dehydratase; CobA – uroporphyrinogen-III C-methyltransferase; CysG – trifunctional enzyme of sirohaem synthesis (see main text); CysK – cysteine synthase A; FolD – bifunctional methylenetetrahydrofolate dehydrogenase (NADP+) / cyclohydrolase; G5K – glutamate 5-kinase; GSD – glutamate semialdehyde dehydrogenase; MTA – 5'-methylthioadenosine; MTFMT – Met-tRNA formyltransferase; OAT – ornithine--oxo-acid transaminase; ODC – ornithine decarboxylase; PSAT – phosphoserine aminotransferase; PSP – phosphoserine phosphatase; (dc)SAM – (decarboxy-)S-adenosylmethionine; SAMD – SAM decarboxylase; SHMT – serine hydroxymethyltransferase; SpS – spermidine/spermin synthase; (f-/m-)THF – (10-formyl-/5,10-methylene-)tetrahydrofolate.

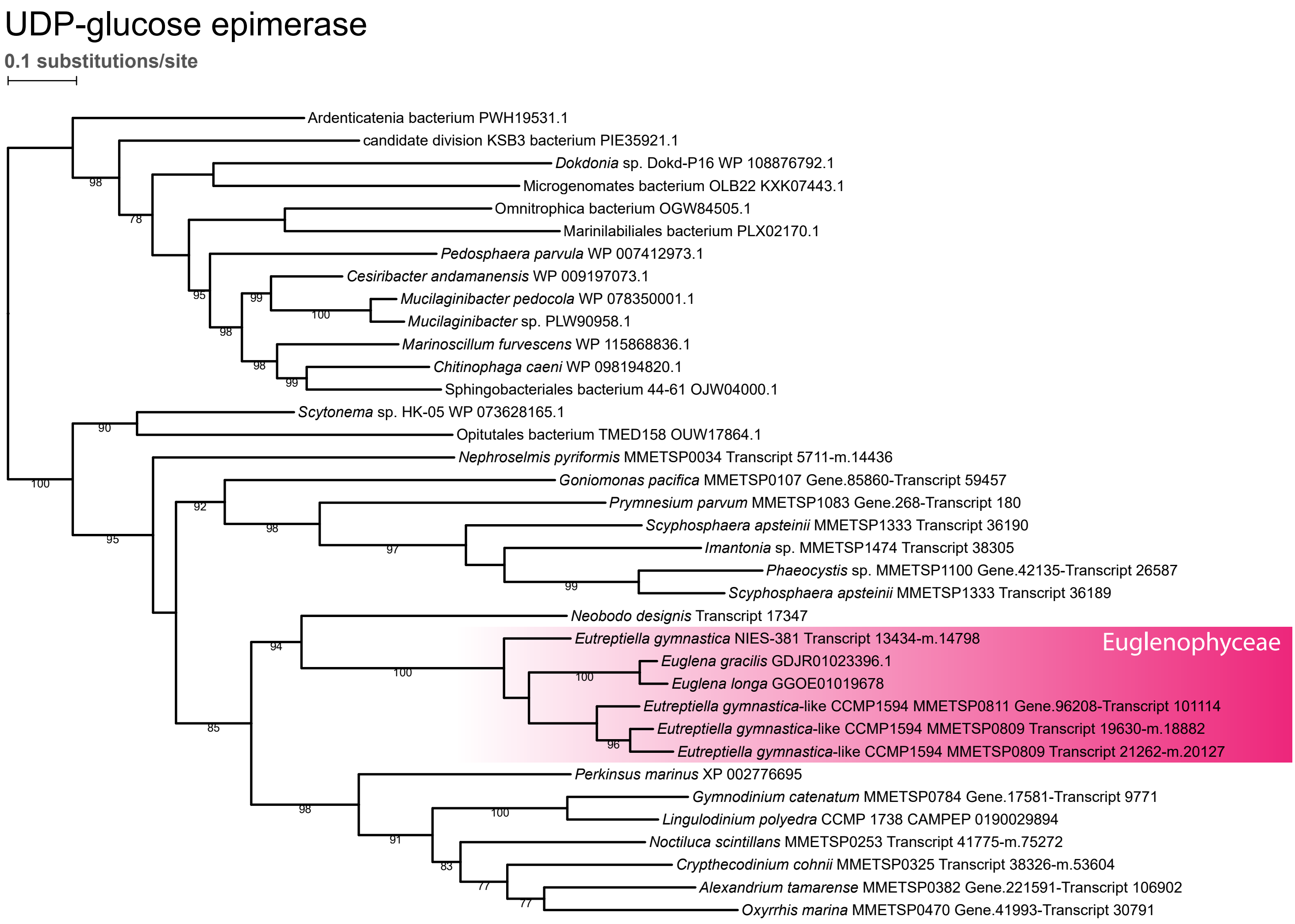


**Figure S5: Inferred phylogeny of the putative plastid UDP-glucose epimerase.** The maximum-likelihood tree was inferred with IQ-TREE using the best-fitting substitution model and ultra-fast bootstrapping. The UFboot support values are indicated at branches when higher than 75%. Colour of the Euglenophyceae clade reflects possible ancestral eukaryotic origin (deep pink).

**triose-phosphate transporters**

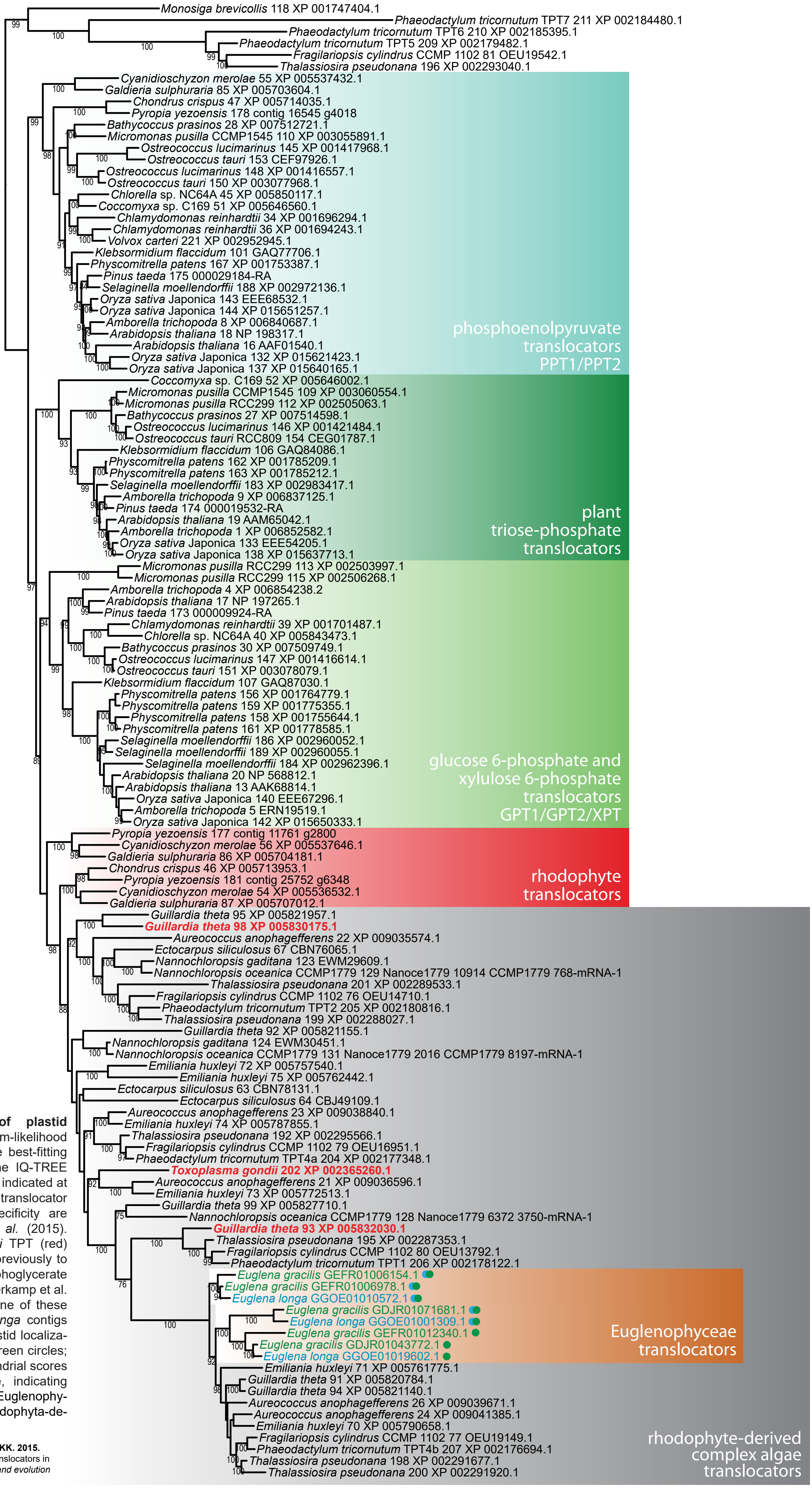


**Figure S6: Inferred phylogeny of plastid phosphate translocators.** The maximum-likelihood tree was inferred with RAxML using the best-fitting substitution model as determined by the IQ-TREE software. The UFboot support values are indicated at branches when higher than 75%. Plant translocator families with determined substrate specificity are marked. Dataset taken from Moog *et al.* (2015). *Guillardia theta* and *Toxoplasma gondii* TPT (red) substrate specificities were determined previously to be DHAP and triose-phosphate/3-phosphoglycerate /phosphoenolpyruvate, respectively (Haferkamp et al. 2006; Brooks et al. 2010). However, none of these translocators clusters closely with *E. longa* contigs (blue) or *E. gracilis* (green). Putative plastid localizations of these sequences is marked by green circles; sequences with high plastid and mitochondrial scores are marked by green and blue circle, indicating possible dual targeting. Colour of the Euglenophyceae clade reflects possible origin in Rhodophyta-derived complex algae (ochre).

**Moog D, Rensing SA, Archibald JM, Maier UG, Ullrich KK. 2015.** Localization and evolution of putative triose phosphate translocators in the diatom *Phaeodactylum tricornutum*. *Genome biology and evolution* **7**: 2955-2969.

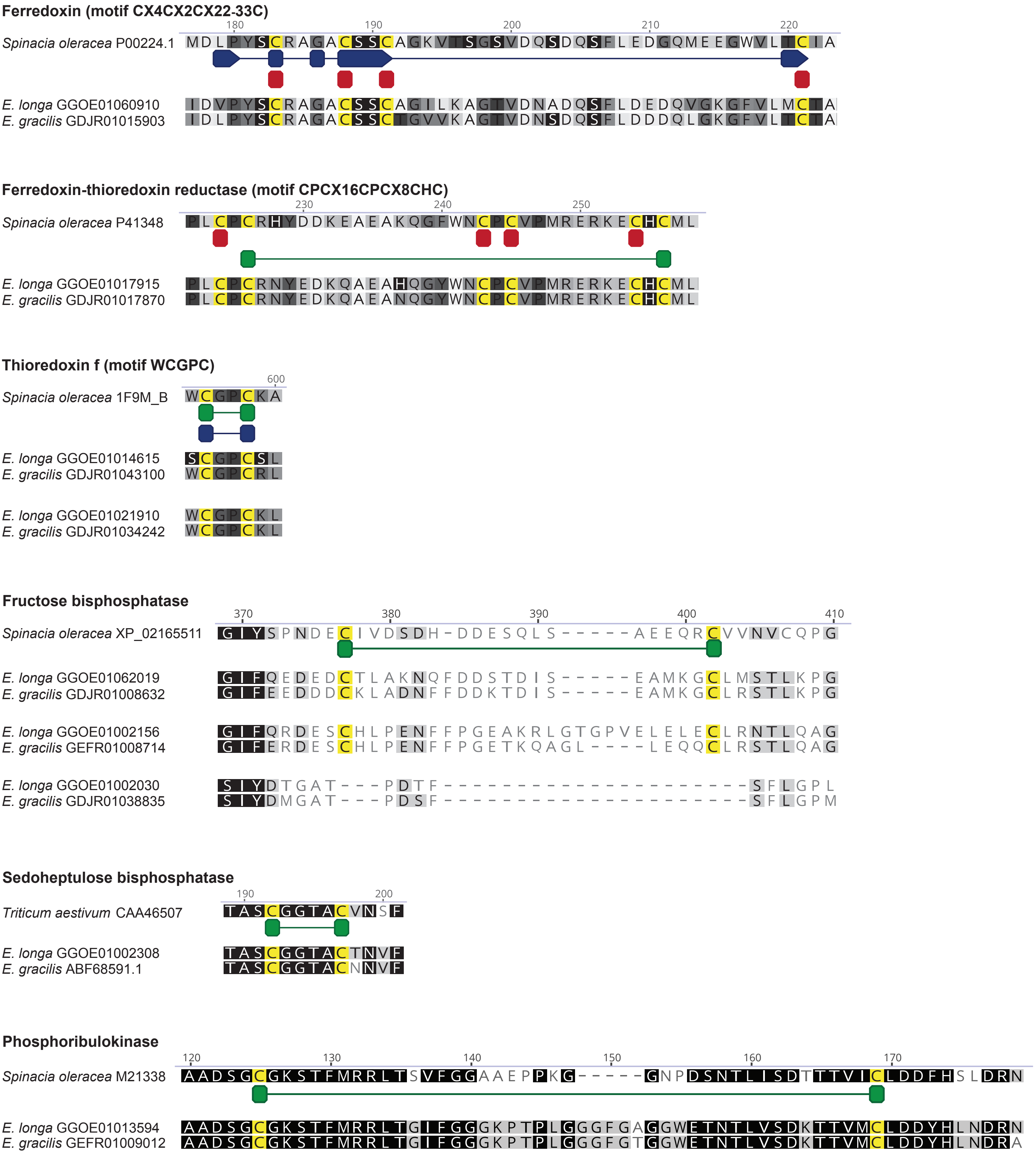


**Figure S7: Conserved cysteine motifs in ferredoxin/thioredoxin system and Calvin-Benson cycle enzymes of *E. longa* and *E. gracilis*.** Shown are alignments of *E. longa* and *E. gracilis* plastid-localized homologs with reference sequences from *Spinacia oleracea*. Connected blue boxes - catalytic residues; red boxes - iron/sulfur cluster binding site; green boxes - redox active cysteine bonds.
