## Supplementary Data - Newick trees for "The cryptic plastid of *Euglena longa* defines a new type of non-photosynthetic plastid organelles"

**Fructose bisphosphate aldolase, dataset listed in Supplementary Table S13:**

((Drosophila melanogaster1:0.12675,Drosophila melanogaster2:0.34855):0.28522,(((Lotharella sp. CCMP6222:0.51769,((Chlorella variabilis1:0.23372,(((((((Arabidopsis thaliana2:0.00862,Arabidopsis thaliana3:0.03349)100:0.03534,Arabidopsis thaliana7:0.05720)97:0.02781,((Oryza sativa Japonica2:0.13246,Oryza sativa Japonica 6:0.03642)100:0.04124,Oryza sativa Japonica4:0.12334)91:0.01251)100:0.03893,(Selaginella moellendorffii2:0.10974,Selaginella moellendorffii3:0.23528)82:0.02889)81:0.02775,((Arabidopsis thaliana4:0.12856,Arabidopsis thaliana9:0.01371)100:0.09255,Oryza sativa Japonica3:0.06824)98:0.03459)91:0.03104,(Physcomitrella patens2:0.07516,Physcomitrella patens4:0.04547)100:0.18773)93:0.04574,Physcomitrella patens5:0.12477)97:0.05448)51:0.03765,Coccomyxa subellipsoidea3:0.20150)67:0.02054)47:0.02450,((((Dictyostelium discoideum:0.28215,((Guillardia theta1:0.11331,Rhodomonas salina2:0.25097)100:0.15052,Cyanidioschyzon merolae1:0.24437)87:0.06092)63:0.02869,(Galdieria sulphuraria2:0.14243,Porphyridium aerugineum2:0.22074)100:0.10060)37:0.02192,(Cryptomonas paramecium1:0.11987,(Guillardia theta2:0.45057,Rhodomonas salina1:0.12742)98:0.07326)100:0.16735)68:0.04526,((((((Neospora caninum1:0.01930,Toxoplasma gondii1:0.02540)100:0.06336,(Neospora caninum2:0.07014,Toxoplasma gondii2:0.00270)99:0.04110)100:0.15122,((Perkinsus marinus1:0.13447,Perkinsus marinus2:0.07994)69:0.03943,Perkinsus marinus3:0.10738)100:0.12979)57:0.04532,((Paramecium tetraurelia1:0.01505,Paramecium tetraurelia2:0.03012)100:0.10633,Tetrahymena thermophila:0.17386)100:0.25161)65:0.04456,(((((Bathycoccus prasinos1:0.32307,(((((Chlamydomonas reinhardtii2:0.02206,Volvox carteri f. nagariensis1:0.02769)100:0.06080,Dunaliella tertiolecta:0.11421)100:0.02248,Polytomella parva:0.17973)99:0.02442,Coccomyxa subellipsoidea1:0.11107)52:0.02989,Chlorella variabilis2:0.10099)100:0.02185)100:0.06239,(((Arabidopsis thaliana1:0.05963,Oryza sativa Japonica1:0.04591)99:0.05876,((((Arabidopsis thaliana5:0.00000,Arabidopsis thaliana8:0.00000)100:0.02248,Arabidopsis thaliana6:0.03123)100:0.03730,Oryza sativa Japonica5:0.04723)100:0.04621,Selaginella moellendorffii1:0.09996)97:0.02541)86:0.01798,((Physcomitrella patens1:0.01807,Physcomitrella patens8:0.01408)100:0.04537,(Physcomitrella patens6:0.02945,Physcomitrella patens7:0.01123)100:0.03545)94:0.02016)98:0.06793)100:0.26753,((Cryptomonas paramecium2:0.19993,((Prochlorococcus marinus:0.34895,Ectocarpus siliculosus2:0.16076)96:0.06276,((Chondrus crispus1:0.09026,(Porphyridium aerugineum1:0.05400,Rhodella maculata1:0.16249)100:0.05906)100:0.04030,(Cyanidioschyzon merolae2:0.21968,Galdieria sulphuraria1:0.09328)97:0.03033)99:0.04922)98:0.02174)96:0.07516,Phaeodactylum tricornutum:0.46689)98:0.05001)89:0.05467,((((((((Chlamydomonas reinhardtii1:0.03675,Volvox carteri f. nagariensis2:0.04117)100:0.04208,Dunaliella tertiolecta1:0.08118)99:0.04019,Coccomyxa subellipsoidea2:0.13049)98:0.04690,Nannochloropsis gaditana:0.21287)45:0.02373,Physcomitrella patens3:0.13292)100:0.14302,Chondrus crispus2:0.30903)100:0.17830,(((Chlamydomonas reinhardtii:0.14697,Volvox carteri f. nagariensis3:0.05660)100:0.35719,Ectocarpus siliculosus3:0.62354)34:0.04741,Chondrus crispus3:0.66344)95:0.09267)47:0.05311,(Euglena gracilis3 PT:0.03990,Euglena longa1 PT:0.02703)100:0.32738)55:0.05155)56:0.03886,(Cyanoptyche gloeocystis:0.27354,((Cyanoptyche gloeocystis2:0.12511,Cyanoptyche gloeocystis3:0.12058)97:0.05853,Gloeochaete wittrockiana2:0.21291)95:0.05016)97:0.04898)41:0.01527)62:0.04948,(Micromonas pusilla2:0.27125,(Ostreococcus tauri1:0.42830,(((Diplonema papillatum:0.29419,(Leptomonas pyrrhocoris:0.09482,Trypanosoma brucei:0.09806)100:0.21065)100:0.38482,Naegleria gruberi:0.49149)91:0.03315,(Nitrobacter hamburgensis:0.39365,Cyanothece sp. PCC 7425:0.47925)100:0.22485)92:0.07354)78:0.04153)87:0.08428)52:0.01731)42:0.01591)70:0.04962,(Lotharella sp. CCMP6223:0.41771,(Gloeochaete wittrockiana:0.30837,(((Lotharella sp. CCMP6221:0.14047,(((Bathycoccus prasinos2:0.09508,Ostreococcus tauri2:0.10580)100:0.07718,Micromonas pusilla1:0.11592)98:0.07090,((Pyramimonas amyliferam:0.16575,Rhodella maculata2:0.50181)96:0.03964,((Euglena gracilis1 PT:0.02104,Euglena longa3 PT:0.07741)100:0.08905,(Eutreptiella gymnastica-like CCMP1594:0.09637,Eutreptiella gymnastica NIES-381:0.05219)99:0.04858)88:0.05169)92:0.03397)100:0.10271)100:0.07412,(Guillardia theta3:0.15680,Ectocarpus siliculosus:0.15417)100:0.10262)100:0.41713,(Euglena gracilis2:0.06640,Euglena longa2:0.10176)100:0.21233)100:0.37878)92:0.05354)52:0.04675):0.01501)100;

**Fructose bisphosphatase, dataset listed in Supplementary Table S14:**

((((((((((((((((((('Eutreptiella gymnastica NIES-381 2':0.15887699999999993,'Eutreptiella gymnastica NIES-381 3':0.10662800000000017)99:0.042946999999999846,'Eutreptiella gymnastica NIES-381 1':0.12924300000000022)95:0.049059000000000186,'Eutreptiella gymnastica-like CCMP1594 1':0.04077900000000012)96:0.04304100000000011,'Euglena longa 7 PT':0.019550999999999874)65:0.006728999999999985,'Euglena gracilis 1 PT':0.018918000000000212)96:0.07441999999999993,('Euglena gracilis 3 PT':0.025993000000000155,'Euglena longa 3 PT':0.0657359999999998)100:0.11893500000000001)99:0.1182120000000002,'Emiliania huxleyi2':0.411127)99:0.06207100000000043,('Phaeodactylum tricornutum1':0.09981600000000013,'Phaeodactylum tricornutum4':0.0826570000000002)100:0.19954300000000025)99:0.0630799999999998,((('Guillardia theta':0.042222000000000204,'Rhodomonas salina2':0.06005899999999986)82:0.04644199999999987,'Cryptomonas paramecium':0.09594700000000023)100:0.15610800000000014,'Emiliania huxleyi3':0.16472599999999993)100:0.06627699999999992)99:0.05322099999999974,(('Phaeodactylum tricornutum2':0.14657799999999988,'Thalassiosira pseudonana2':0.1809210000000001)100:0.3103030000000002,'Ectocarpus siliculosus2':0.6496560000000002)100:0.38211799999999974)80:0.06268700000000038,((('Phaeodactylum tricornutum5':0.32893300000000014,'Thalassiosira pseudonana1':0.3466070000000001)100:0.43864400000000003,'Ectocarpus siliculosus1':0.14325300000000007)69:0.005465000000000053,'Aureococcus anophagefferens1':0.2012900000000002)65:0.02129700000000012)66:0.040765999999999636,'Lotharella sp. CCMP6223':0.28142699999999987)60:0.03072800000000031,(((('Porphyridium aerugineum3':0.305971,'Porphyridium aerugineum1':0.14819300000000002)94:0.0709850000000003,'Chondrus crispus2':0.06537800000000038)82:0.014049999999999674,('Rhodella maculata1':0.02111900000000011,'Rhodella maculata2':0.0032840000000002867)100:0.16647699999999999)85:0.03274599999999994,(('Galdieria sulphuraria':0.31458200000000014,'Galdieria sulphuraria2':0.1218309999999998)96:0.028289999999999704,'Cyanidioschyzon merolae':0.3561599999999996)96:0.08015000000000017)96:0.08085999999999993)78:0.105715,((((((((('Chlamydomonas reinhardtii':0.017441999999999958,'Volvox carteri f. nagariensis':0.026727000000000167)100:0.034191000000000304,'Polytomella parva':0.11968799999999957)100:0.03473999999999977,'Dunaliella tertiolecta2':0.1565669999999999)100:0.04584400000000022,'Chlorella variabilis2':0.29544599999999965)97:0.03007900000000019,'Coccomyxa subellipsoidea3':0.12199900000000019)96:0.04178499999999952,'Pyramimonas parkeae':0.367289)17:0.000002,((('Bathycoccus prasinos':0.21864099999999986,'Micromonas pusilla3':0.19259799999999982)100:0.42585300000000004,'Micromonas pusilla':0.12205399999999988)66:0.040863000000000316,('Bathycoccus prasinos2':0.16168499999999986,'Ostreococcus tauri':0.040134000000000114)100:0.08139200000000013)72:0.03501099999999946)84:0.04786900000000038,((('Physcomitrella patens2':0.023608000000000295,'Physcomitrella patens3':0.01407500000000006)100:0.05701299999999998,'Selaginella moellendorffii3':0.07462900000000028)76:0.02430500000000002,('Arabidopsis thaliana2':0.029383999999999855,'Oryza sativa2':0.077909)78:0.02223200000000025)99:0.050060000000000215)99:0.09970799999999969,((('Coccomyxa subellipsoidea2':0.20696599999999998,'Dunaliella tertiolecta':0.3928449999999999)41:0.07094000000000023,'Porphyridium aerugineum4':0.5125489999999999)99:0.14308500000000013,(('Euglena gracilis 6 PT':0.15140999999999982,'Euglena longa 2 PT':0.12530300000000016)100:0.31376800000000005,'Lotharella sp. CCMP622':0.449319)100:0.21383300000000016)100:0.270197)84:0.0880209999999999)50:0.018107000000000095,(('Gloeochaete wittrockiana3':0.3962979999999998,'Gloeochaete wittrockiana2':0.08327300000000015)52:0.053160000000000096,('Cyanoptyche gloeocystis1':0.04277799999999976,'Cyanoptyche gloeocystis2':0.02212899999999962)100:0.16056000000000026)58:0.03960000000000008)54:0.05656999999999979,(('Rhodomonas salina3':0.13154399999999988,'Guillardia theta2':0.04964000000000013)95:0.03974999999999973,'Cryptomonas paramecium2':0.14526199999999978)100:0.427508)97:0.138725,(((((((((((sbpChondrus:0.31892299999999985,sbpGaldieria2:0.19284999999999997)97:0.11605200000000027,sbpCyanidioschyzon:0.5246210000000002)85:0.07597299999999985,(sbpEmiliania1:1.1581300000000003,sbpGaldieria:0.47376600000000035)82:0.07718299999999978)96:0.14341000000000026,(((sbpChlamydomonas:0.014845999999999915,sbpVolvox:0.017534999999999634)100:0.09454300000000027,sbpCoccomyxa:0.1350659999999997)97:0.065137,sbpArabidopsis:0.21798099999999998)100:0.28804700000000016)78:0.167767,sbpNannochloropsis:0.9085800000000002)98:0.17170599999999991,(((sbpEmiliania2:0.4229890000000003,sbpEmiliania3:0.15147900000000014)97:0.20577500000000004,sbpOstreococcus:0.1581520000000003)100:0.311763,sbpToxoplasma:0.6328290000000001)100:0.3234079999999997)95:0.10732700000000017,(sbpTrypanosoma:0.7562679999999999,sbpCyanidioschyzon2:0.6635750000000002)84:0.08363299999999985)100:0.7751490000000003,'Diplonema papillatum':1.2187039999999998)89:0.27170099999999975,('Trypanosoma brucei':0.233854,'Leishmania major':0.17471000000000014)100:0.275369)58:0.11631299999999989,((('Chondrus crispus':0.0860369999999997,'Galdieria sulphuraria3':0.2141820000000001)100:0.051741999999999955,'Cyanidioschyzon merolae2':0.12914799999999982)91:0.04824000000000028,'Porphyridium aerugineum2':0.24015699999999995)100:0.09526799999999991)60:0.03988099999999983,('Gloeochaete wittrockiana':0.37178799999999956,'Dictyostelium discoideum':0.3685510000000001)98:0.16424700000000048)61:0.08534000000000042)56:0.033564999999999845,((((((((((((('Euglena longa 4':0.11075200000000018,'Euglena longa 5':0.000002)100:0.02280300000000013,'Euglena gracilis 4':0.007054999999999811)100:0.0437470000000002,'Eutreptiella gymnastica-like CCMP1594 2':0.09976299999999982)100:0.0592649999999999,'Eutreptiella gymnastica-like CCMP1594 3':0.0868190000000002)63:0.021977000000000135,('Emiliania huxleyi':0.19615699999999991,'Eutreptiella gymnastica-like CCMP1594 5':0.06357299999999988)92:0.08167200000000019)31:0.000002,('Euglena longa 6':0.010639999999999983,'Euglena gracilis 2':0.05032300000000012)100:0.07950200000000018)91:0.11016400000000015,(('Skeletonema marinoi':0.021389000000000102,'Thalassiosira pseudonana3':0.050044999999999895)100:0.11949299999999985,'Phaeodactylum tricornutum3':0.06429499999999999)100:0.2486830000000002)97:0.08065999999999995,('Lotharella sp. CCMP6222':0.46784199999999965,'Aureococcus anophagefferens2':0.2100259999999996)65:0.1219800000000002)84:0.1584080000000001,('Phytophtora ramorum':0.13333200000000023,'Phytophtora ramorum2':0.049129999999999896)100:0.1724899999999998)100:0.1021040000000002,'Monosiga brevicollis':0.42358300000000026)89:0.09610099999999999,(((((((('Oryza sativa':0.08854599999999957,'Arabidopsis thaliana':0.030199000000000087)99:0.027324000000000126,'Physcomitrella patens1':0.09769300000000003)97:0.0069509999999999295,('Selaginella moellendorffii1':0.016596999999999973,'Selaginella moellendorffii4':0.000002)100:0.04638600000000004)100:0.050035000000000274,'Selaginella moellendorffii2':0.05751100000000031)100:0.141972,'Coccomyxa subellipsoidea':0.3711750000000005)74:0.030092999999999925,('Ostreococcus tauri2':0.9602250000000003,'Micromonas pusilla2':0.36929900000000027)62:0.13163400000000003)58:0.02289799999999964,'Chlorella variabilis':0.39515900000000004)39:0.014427000000000412,(('Cryptococcus neoformans':0.0742149999999997,'Laccaria bicolor':0.0967640000000003)100:0.14975899999999998,'Aspergillus fumigatus':0.5421209999999999)65:0.12892800000000015)56:0.03264099999999992)63:0.04488799999999982,'Galdieria sulphuraria4':0.8071260000000002)50:0.022387999999999852,(((('Guillardia theta3':0.08943399999999979,'Rhodomonas salina1':0.2186720000000002)100:0.38520999999999983,'Tetrahymena thermophila':0.3070789999999999)89:0.031181000000000125,'Drosophila melanogaster':0.427613)84:0.06958999999999982,'Acanthamoeba castellanii':0.36736900000000006)75:0.03323300000000007)59:0.03770500000000032):0.07927699999999982,((((((('Euglena gracilis 5':0.017386000000000124,'Euglena longa 1':0.03818700000000019)100:0.10832400000000009,'Eutreptiella gymnastica-like CCMP1594 4':0.10041600000000006)100:0.3407300000000002,'Cupriavidus necator':0.32853699999999986)81:0.03979299999999997,'Magnetospirillum magneticum':0.21201099999999995)86:0.035679000000000016,((('Cupriavidus necator2':0.03828200000000015,'Ralstonia solanacearum':0.07962099999999994)100:0.03723799999999988,'Verminephrobacter eiseniae':0.17688199999999998)98:0.04211099999999979,'Burkholderia cenocepacia':0.1798329999999999)100:0.1815190000000002)100:0.3347640000000003,((((('Synechococcus sp. PCC 7335':0.1960449999999998,'Thermosynechococcus elongatus':0.13190800000000014)66:0.05911799999999978,'Cyanothece sp. PCC 7425':0.09604100000000004)66:0.03500099999999984,'Lyngbya sp. PCC 8106':0.1278060000000001)65:0.058803999999999856,('Anabaena variabilis':0.013650000000000162,'Nodularia spumigena':0.017787999999999915)100:0.03747100000000003)100:0.34835800000000017,('Cyanoptyche gloeocystis3':1.793238,'Synechococcus sp. PCC 73352':0.5530400000000002)76:0.20573500000000022)83:0.18491500000000016)58:0.09702699999999975,((('Neospora caninum':0.006352000000000135,'Toxoplasma gondii':0.000002)99:0.016659999999999897,'Eimeria tenella':0.10695299999999985)100:0.21356399999999987,('Perkinsus marinus':0.44943,'Yersinia pestis':0.24291000000000018)45:0.042714999999999836)100:0.31011500000000014)76:0.07927700000000026);

**Glyceraldehyde phosphate dehydrogenase, dataset listed in Supplementary Table S15:**

((((((((((((((((((('Thalassiosira pseudonana':0.000003,'Thalassiosira pseudonana2':0.000003)100:0.05572200000000005,'Skeletonema marinoi':0.04011588780000008)81:0.028784000000000032,'Thalassiosira pseudonana3':0.05250136830000007)100:0.08428599999999997,(('Durinskia baltica2':0.08112541940000007,'Phaeodactylum tricornutum':0.0671700972999999)80:0.035804999999999976,'Phaeodactylum tricornutum2':0.0065558436999999525)80:0.03768499999999997)100:0.06405400000000006,(('Ochromonas sp. CCMP1393':0.0463296809,'Durinskia baltica1':0.07292459759999992)100:0.035978000000000065,'Dinobryon sp. UTEXLB2267':0.038816452100000065)100:0.128927)31:0.024255999999999944,'Aureococcus anophagefferens':0.13696430729999998)29:0.013548000000000004,(('Ectocarpus siliculosus':0.1322594734,'Ectocarpus siliculosus2':0.11794454099999996)78:0.04810499999999995,'Phytophtora ramorum':0.17138284429999995)70:0.04318999999999995)78:0.09727700000000006,(((('Laccaria bicolor':0.04140342149999998,'Laccaria bicolor2':0.06589186759999999)100:0.12331400000000003,'Cryptococcus neoformans':0.09043171679999995)88:0.04127400000000003,('Aspergillus fumigatus':0.11481301919999998,'Aspergillus fumigatus2':0.20973984089999997)100:0.07048600000000005)77:0.03251999999999999,'Drosophila melanogaster':0.195860826)60:0.008301000000000003)49:0.01611600000000002,(((('Arabidopsis thaliana2':0.019468537399999986,'Arabidopsis thaliana3':0.02975680709999995)100:0.060078999999999994,('Oryza sativa':0.011736301699999951,'Oryza sativa6':0.035320910799999994)97:0.018279000000000045)100:0.08600399999999997,'Selaginella moellendorffii':0.10666085719999996)98:0.052100000000000035,('Physcomitrella patens8':0.000002,'Physcomitrella patens9':0.013879531700000003)100:0.10204400000000002)89:0.02954100000000004)61:0.02358899999999997,'Monosiga brevicollis':0.25061763319999997)47:0.009552000000000005,(((((('Perkinsus marinus':0.0041020647999999715,'Perkinsus marinus4':0.032397326000000004)84:0.017974000000000045,'Perkinsus marinus2':0.04490093849999999)100:0.09651900000000002,'Perkinsus marinus3':0.10893468289999997)99:0.031046000000000018,'Neospora caninum':0.2983331683999999)100:0.06768099999999999,(('Chattonella subsalsa':0.1003730745000001,'Heterosigma akashiwo':0.038651048700000024)100:0.13237,'Phytophtora ramorum2':0.17053460369999995)98:0.06147800000000003)99:0.02527299999999999,'Entamoeba histolytica':0.2591058935)76:0.020236000000000032)44:0.017009000000000052,((((((('Pyramimonas amylifera2':0.13729370909999994,'Pyramimonas parkeae2':0.07635991210000004)100:0.07026900000000003,'Chlorella variabilis':0.14560066159999996)75:0.022070000000000034,'Coccomyxa subellipsoidea':0.0862553533)52:0.04236099999999998,(('Chlorella variabilis2':0.18216206359999998,'Coccomyxa subellipsoidea3':0.1623986851)61:0.049634999999999985,'Chlamydomonas reinhardtii2':0.2216520634)56:0.021974999999999967)84:0.04506999999999994,((('Chondrus crispus2':0.14421536840000004,'Cyanidioschyzon merolae2':0.2690183059)72:0.04901500000000003,('Galdieria sulphuraria2':0.12111790700000002,'Rhodella maculata':0.14819737060000004)80:0.029518000000000044)44:0.027436000000000016,'Porphyridium aerugineum2':0.07818751499999999)90:0.04614099999999999)67:0.024685999999999986,((((('Oryza sativa2':0.01746840670000005,'Oryza sativa7':0.08668837330000001)100:0.015225999999999962,'Oryza sativa5':0.004443217299999969)100:0.036082999999999976,'Arabidopsis thaliana5':0.04724303129999996)99:0.05437499999999995,'Selaginella moellendorffii2':0.11173577430000003)99:0.04363600000000001,('Physcomitrella patens':0.033514984199999986,'Physcomitrella patens10':0.01504179579999998)100:0.05415700000000001)99:0.06679000000000002)13:0.018861000000000017,(('Cyanoptyche gloeocystis':0.11676831200000004,'Gloeochaete wittrockiana2':0.09085054790000002)100:0.07971499999999998,'Gloeochaete wittrockiana3':0.15207961969999995)94:0.06635599999999997)9:0.000002)38:0.008677999999999964,((('Dictyostelium discoideum':0.2082410361,'Bathycoccus prasinos3':0.17762288120000003)87:0.04863099999999998,('Guillardia theta':0.03638453399999997,'Rhodomonas salina':0.09277264070000002)100:0.16644199999999998)73:0.03917700000000002,'Micromonas pusilla3':0.22784358380000003)59:0.018940999999999986)65:0.05896800000000002,'Lotharella sp. CCMP622':0.21308649179999994)48:0.018375999999999948,('Porphyromonas gingivalis':0.13972456099999997,'Trypanosoma brucei':0.18338154409999996)97:0.08063299999999995)21:0.008009999999999962,((('Eutreptiella gymnastica-like CCMP1594 1':0.08077856910000003,'Eutreptiella gymnastica NIES-381 3':0.12322109589999997)98:0.05020999999999998,'Euglena longa 2':0.4063773786)100:0.11888200000000004,'Naegleria gruberi':0.21203204669999998)71:0.09956399999999999)35:0.023805999999999994,'Bigelowiella natans':0.23876163839999998)99:0.131231,(('Euglena gracilis 2':0.04038767859999992,'Euglena longa 3':0.02053471809999996)100:0.12470400000000015,('Euglena gracilis 3':0.029687081200000076,'Euglena longa 1':0.012545785899999995)97:0.019628999999999897)100:0.3753900000000001):0.15039899999999995,((((((((((((('Physcomitrella patens4':0.000003,'Physcomitrella patens5':0.000003)100:0.008507000000000042,'Physcomitrella patens3':0.022132972399999895)99:0.00468500000000005,'Physcomitrella patens2':0.005181038099999924)100:0.01252399999999998,('Physcomitrella patens6':0.000003,'Physcomitrella patens7':0.000003)96:0.000003)100:0.03264899999999993,(('Arabidopsis thaliana6':0.037489140299999946,'Oryza sativa4':0.04710140049999989)100:0.019895999999999914,'Arabidopsis thaliana4':0.003416211900000077)100:0.04578400000000005)96:0.022407999999999983,'Selaginella moellendorffii3':0.047152670800000074)100:0.04048199999999991,((('Chlamydomonas reinhardtii':0.042694608399999945,'Volvox carteri f. nagariensis':0.004245143799999962)100:0.030043000000000042,('Dunaliella tertiolecta':0.000002,'Noctiluca scintillans2':0.000003)100:0.09551799999999999)100:0.028362999999999916,'Coccomyxa subellipsoidea2':0.09053610860000005)54:0.027336999999999945)86:0.02639900000000006,(((('Bathycoccus prasinos':0.04468919539999994,'Ostreococcus tauri':0.014721661899999905)85:0.01462399999999997,'Micromonas pusilla2':0.016580099700000073)100:0.08660599999999996,(('Bathycoccus prasinos2':0.06038359000000004,'Ostreococcus tauri2':0.08210435929999993)97:0.030539000000000094,'Micromonas pusilla':0.09036646820000005)100:0.0479989999999999)38:0.022134000000000098,((('Arabidopsis thaliana':0.051433444800000006,'Oryza sativa3':0.039344505499999904)99:0.02736699999999992,'Selaginella moellendorffii4':0.048046216799999986)98:0.021598999999999924,'Pyramimonas amylifera3':0.15114099889999988)100:0.03976399999999991)83:0.011732999999999993)89:0.0367559999999999,((('Cyanoptyche gloeocystis2':0.000003,'Cyanoptyche gloeocystis3':0.004435895300000103)100:0.09308300000000003,'Gloeochaete wittrockiana':0.05053279339999994)90:0.03452199999999994,'Cyanophora paradoxa':0.051239912800000065)100:0.0750050000000001)74:0.015185000000000004,(((((('Porphyridium aerugineum':0.13151494789999996,'Porphyra purpurea':0.14134259139999994)90:0.025023000000000017,'Chondrus crispus':0.10295799690000007)62:0.019729000000000108,'Galdieria sulphuraria':0.1504808032)90:0.02665699999999993,'Rhodella maculata2':0.1257134324)99:0.06085099999999999,'Cyanidioschyzon merolae':0.19214032669999992)78:0.051378000000000035,(('Pyramimonas parkeae':0.029391393999999904,'Pyramimonas parkeae3':0.030284812399999916)73:0.01849400000000001,'Pyramimonas amylifera':0.05205733080000008)100:0.09651599999999994)70:0.018070999999999948)81:0.05328500000000003,((((('Anabaena variabilis':0.03364628230000011,'Nodularia spumigena':0.03409136359999998)100:0.10837999999999992,('Cyanothece sp. PCC 7425':0.03257068919999995,'Thermosynechococcus elongatus':0.0997534391999999)100:0.09965399999999991)85:0.04984199999999994,'Synechococcus sp. PCC 7335':0.20837697519999998)68:0.0158910000000001,(('Paulinella chromatophora':0.09827099419999996,'Prochlorococcus marinus':0.15309134590000006)100:0.18735999999999997,'Crocosphaera watsonii':0.07130313479999995)78:0.030980000000000008)91:0.055169000000000024,'Lyngbya sp. PCC 8106':0.0870566948)100:0.112545)99:0.06402600000000003,(((((('Eutreptiella gymnastica NIES-381 1':0.030677989399999994,'Eutreptiella gymnastica NIES-381 2':0.040970434799999955)100:0.012388999999999983,'Eutreptiella gymnastica NIES-381 4':0.02601348119999991)54:0.011646999999999963,'Eutreptiella gymnastica NIES-381 5':0.02561392320000011)100:0.07001100000000005,('Euglena gracilis 1PT':0.06237797239999998,'Eutreptiella gymnastica-like CCMP1594 2':0.06790787719999991)100:0.04058800000000007)100:0.06228,('Crypthecodinium cohniiP':0.13475968640000002,'Noctiluca scintillans1':0.11278989489999991)100:0.09948199999999996)100:0.06492500000000001,'Oxyrrhis marina':0.5338145014)100:0.07393399999999994)100:0.16216799999999998,'Bacillus anthracis':0.2872151165)100:0.15039900000000006);

**Phosphoglycerate kinase, dataset listed in Supplementary Table S16:**

(((((((((((((((((((('Durinskia baltica':0.030665879600000112,'Kryptoperidinium foliaceum4':0.021590375099999948)100:0.03726800000000008,'Phaeodactylum tricornutum':0.04129386440000005)100:0.028804000000000052,'Thalassiosira pseudonana':0.026647592800000064)99:0.033822999999999936,'Skeletonema marinoi':0.04484207270000007)95:0.04665600000000003,'Noctiluca scintillans2':0.05567403570000007)100:0.15990900000000008,'Aureococcus anophagefferens':0.18968105460000007)98:0.032010000000000094,(('Durinskia baltica3':0.09322199819999999,'Lingulodinium polyedra2':0.055324120599999915)99:0.016192999999999902,'Kryptoperidinium foliaceum3':0.0883137987)100:0.22623700000000002)86:0.04282500000000011,'Ectocarpus siliculosus2':0.0861992775)80:0.014475999999999933,'Vaucheria litorea':0.12835901979999997)89:0.023773999999999962,(('Cryptomonas paramecium2':0.1459782181,'Guillardia theta2':0.04819933550000011)100:0.008912999999999949,('Guillardia theta1':0.05201570420000001,'Rhodomonas salina2':0.06022198259999989)100:0.03216400000000008)100:0.08990000000000009)89:0.03187300000000004,('Chattonella subsalsa':0.027456385000000028,'Heterosigma akashiwo':0.03383132900000008)100:0.050875000000000004)85:0.023676999999999948,((((((('Dinobryon sp. UTEXLB22671':0.041996847200000076,'Ochromonas sp. CCMP1393':0.09077862010000004)99:0.05384699999999998,('Dinobryon sp. UTEXLB22672':0.000003,'Dinobryon sp. UTEXLB22673':0.007244248099999906)100:0.07401900000000006)100:0.10178200000000004,'Ectocarpus siliculosus':0.11503067450000004)69:0.017217000000000038,(('Chondrus crispus':0.059588754600000016,'Porphyra purpurea':0.10509847259999994)99:0.021625999999999923,('Porphyridium aerugineum':0.10060551679999996,'Rhodella maculata':0.08292647129999997)94:0.014796000000000031)99:0.064249)33:0.015867000000000075,('Bigelowiella natans':0.041197753699999895,'Lotharella sp. CCMP622':0.05528038269999991)100:0.092171)75:0.013071999999999973,(('Karenia brevisS2':0.028607059200000062,'Karenia brevisS4':0.047084012200000025)100:0.06429599999999991,('Pleurochrysis carterae2':0.008217336299999989,'Prymnesium parvum Texoma12':0.06730977290000006)100:0.04151199999999999)100:0.043147000000000046)76:0.01374300000000006,('Nannochloropsis gaditana1':0.20772789339999997,'Nannochloropsis gaditana2':0.06545243079999996)99:0.09411999999999998)67:0.01182799999999995)57:0.000002,('Cyanidioschyzon merolae':0.15488740209999996,'Galdieria sulphuraria':0.2254667724999999)99:0.10616599999999998)76:0.04563000000000006,(((((((('Anabaena variabilis':0.016300713399999944,'Nodularia spumigena':0.07253446199999991)100:0.04083700000000001,'Crocosphaera watsonii':0.11661893620000008)93:0.02352299999999996,'Cyanothece sp. PCC 7425':0.08535974940000002)48:0.018291999999999975,('Lyngbya sp. PCC 8106':0.08053120260000002,'Synechococcus sp. PCC 7335':0.12729535600000008)83:0.05152599999999996)46:0.02017000000000002,'Thermosynechococcus elongatus':0.12562924570000011)72:0.03026200000000001,('Paulinella chromatophora':0.06037673890000006,'Prochlorococcus marinus':0.17295040249999993)100:0.08594900000000005)100:0.10901000000000005,((((('Oryza sativa':0.03194477870000001,'Oryza sativa2':0.0220881341000001)100:0.05364600000000008,'Arabidopsis thaliana3':0.06994662760000003)100:0.056486000000000036,'Selaginella moellendorffii2':0.16853129729999994)90:0.011061999999999905,('Arabidopsis thaliana':0.019560816600000086,'Arabidopsis thaliana2':0.01202463400000009)100:0.042596999999999996)96:0.020351999999999926,'Selaginella moellendorffii':0.03642811789999989)100:0.04038300000000006)88:0.041881999999999975,(((('Polytomella parva':0.17647080569999996,'Volvox carteri f. nagariensis':0.0337611914)75:0.007913000000000059,'Chlamydomonas reinhardtii':0.004884838099999955)100:0.03926000000000007,'Dunaliella tertiolecta':0.2180328866000001)96:0.03781600000000007,((('Pyramimonas amylifera':0.0707957292000001,'Pyramimonas parkeae':0.0509621467000001)100:0.07950999999999997,'Chlorella variabilis':0.08060426730000003)87:0.01334499999999994,'Coccomyxa subellipsoidea':0.12443157149999995)86:0.019662999999999986)94:0.024494999999999933)87:0.017152000000000056)78:0.06613100000000005,('Cyanophora paradoxa':0.09488314300000011,'Gloeochaete wittrockiana':0.1289492564999999)91:0.019477000000000078)66:0.019139000000000017,'Cyanoptyche gloeocystis':0.13255010760000008)98:0.058011000000000035,(('Physcomitrella patens':0.05553479760000002,'Physcomitrella patens2':0.07319342130000006)100:0.037717,'Physcomitrella patens3':0.09449717430000004)100:0.12053400000000003)100:0.1848280000000001,(('Bacteroides fragilis':0.057276487700000045,'Porphyromonas gingivalis':0.15187745130000008)100:0.269246,'Flavobacterium columnare':0.36214994080000007)100:0.2386290000000001)73:0.030310999999999755,('Bacillus anthracis':0.3706539753000001,'Listeria monocytogenes':0.5229608721)55:0.051409999999999845)65:0.03026799999999996,(((((((((((('Eutreptiella gymnastica-like CCMP1594 1':0.000003,'Eutreptiella gymnastica-like CCMP1594 4':0.000003)100:0.03838800000000009,'Eutreptiella gymnastica NIES-381 1':0.0687214585)100:0.03155299999999994,('Euglena gracilis 4':0.00768352129999994,'Euglena gracilis 3PT':0.006776149600000059)100:0.0431919999999999)100:0.110908,(((('Guillardia theta3':0.06219247449999998,'Rhodomonas salina':0.05252402190000005)93:0.035813000000000095,'Cryptomonas paramecium':0.07293295090000007)95:0.04507799999999995,'Emiliania huxleyi':0.3682332480000001)74:0.02856600000000009,'Oxyrrhis marina':0.13119874889999994)65:0.017622000000000027)74:0.016378000000000004,('Emiliania huxleyi2':0.015147489799999914,'Emiliania huxleyi3':0.000003)100:0.286138)75:0.02747799999999989,(('Pleurochrysis carterae3':0.1021166402,'Prymnesium parvum Texoma1':0.13728192950000007)90:0.035420999999999925,'Pleurochrysis carterae':0.11463107100000003)65:0.02822199999999997)48:0.022407000000000066,(('Symbiodinium sp. CCMP4211':0.04950776689999992,'Symbiodinium sp. CCMP4214':0.05629302040000006)100:0.07966899999999999,'Durinskia baltica2':0.16912363419999998)94:0.046221999999999985)54:0.035744999999999916,(((((('Karenia brevisS3':0.09562415289999993,'Karenia brevisS6':0.16331370320000005)68:0.0250999999999999,'Karenia brevisS5':0.09719401249999993)71:0.039309999999999956,'Kryptoperidinium foliaceum1':0.054291290899999955)63:0.01527400000000001,('Lingulodinium polyedra1':0.11774873709999989,'Lingulodinium polyedra3':0.02861367700000006)88:0.021309000000000022)73:0.020590000000000108,('Crypthecodinium cohnii':0.10560663850000007,'Noctiluca scintillans1':0.06889416589999997)100:0.14368099999999995)62:0.02168900000000007,('Symbiodinium sp. CCMP4212':0.043648585599999956,'Symbiodinium sp. CCMP4215':0.05037811709999995)83:0.015594000000000108)84:0.05071899999999996)30:0.03252699999999997,(((('Symbiodinium sp. CCMP4213':0.047957427899999905,'Symbiodinium sp. CCMP4216':0.016188470999999982)100:0.039323,'Crypthecodinium cohnii2':0.10194836809999996)99:0.029274000000000022,('Karenia brevisS1':0.04025845639999992,'Karenia brevisS7':0.017914468699999908)100:0.09646700000000008)64:0.010434999999999972,('Kryptoperidinium foliaceum2':0.1163206416,'Kryptoperidinium foliaceum5':0.06715660579999994)97:0.021239000000000008)99:0.03154799999999991)100:0.16598800000000002,('Eutreptiella gymnastica-like CCMP1594 3':0.14380870690000003,'Phytophtora ramorum':0.17555410449999997)100:0.04446400000000006)98:0.09722900000000001,(('Euglena gracilis 2':0.019368422499999927,'Euglena longa 1':0.0936283752)100:0.334284,('Naegleria gruberi':0.46789507539999997,'Laccaria bicolor':0.2058643548000001)79:0.08739699999999995)70:0.022035000000000027)100:0.2986329999999999,((((('Euglena gracilis 1B':0.03190014969999999,'Euglena gracilis 1':0.000003)100:0.02784399999999998,'Euglena longa 2':0.01660041379999999)100:0.052000999999999964,('Eutreptiella gymnastica-like CCMP1594 2':0.11269512339999999,'Eutreptiella gymnastica NIES-381 2':0.08791770850000002)100:0.039795000000000025)100:0.2972159999999999,((('Trypanosoma brucei':0.00746156259999986,'Trypanosoma brucei3':0.16204816099999986)92:0.009740000000000082,'Trypanosoma brucei2':0.012119018599999976)99:0.13055700000000003,'Leishmania major1':0.09884054080000015)100:0.1950860000000001)100:0.1128610000000001,'Leishmania major2':0.8454950784999999)99:0.14687300000000003):0.030268000000000184);

**Phosphoribulokinase, dataset listed in Supplementary Table S17:**

((((((((((('Guillardia theta':0.15596538319999997,'Rhodomonas salina':0.2352551476)98:0.08298499999999998,'Cryptomonas paramecium':0.27971967650000007)100:0.16249599999999997,('Phaeodactylum tricornutum':0.15643176859999997,'Thalassiosira pseudonana':0.15039566930000003)84:0.12344500000000003)92:0.06319699999999995,'Emiliania huxleyi':0.5525154438)50:0.07617700000000005,(((('Euglena gracilis':0.02425356550000002,'Euglena longa':0.05964046940000001)97:0.060953000000000035,'Eutreptiella gymnastica-like CCMP1594':0.06498231160000001)47:0.05077200000000004,'Eutreptiella gymnastica NIES-381':0.05819824149999997)100:0.42819299999999993,'Ectocarpus siliculosus':0.10996973489999995)81:0.06657100000000005)89:0.07881900000000008,(('Bathycoccus prasinos':0.06808705000000004,'Ostreococcus tauri':0.046814661300000004)71:0.011155999999999944,'Micromonas pusilla':0.027853968399999984)100:0.08160900000000004)44:0.03805499999999995,(((('Chlamydomonas reinhardtii':0.02106741550000013,'Volvox carteri f. nagariensis':0.020189970400000035)100:0.02940900000000002,'Dunaliella tertiolecta':0.11402544650000013)93:0.01048899999999997,'Chlorella variabilis':0.07429665480000003)100:0.051775000000000015,((('Pyramimonas parkeae':0.000003,'Pyramimonas parkeae2':0.1606920464)100:0.053921999999999914,'Pyramimonas amylifera':0.03300271619999995)99:0.05607899999999999,'Coccomyxa subellipsoidea':0.08343717299999998)91:0.04861900000000008)30:0.014168999999999987)47:0.026939000000000046,((('Oryza sativa':0.02585452340000005,'Oryza sativa2':0.11100106030000001)86:0.008682000000000079,'Arabidopsis thaliana':0.0419007374)100:0.03394799999999987,(('Selaginella moellendorffii':0.000003,'Selaginella moellendorffii2':0.003870475100000048)100:0.03197399999999995,'Physcomitrella patens':0.04238081260000004)70:0.00789799999999996)97:0.023694000000000104)100:0.136563,(((('Cyanoptyche gloeocystis2':0.000003,'Cyanoptyche gloeocystis3':0.008080745300000025)100:0.055709999999999926,'Cyanoptyche gloeocystis':0.0350837651)99:0.05484,'Gloeochaete wittrockiana':0.08069170530000003)85:0.026403000000000065,'Gloeochaete wittrockiana2':0.38582231990000004)99:0.06507600000000002)76:0.027302999999999966,(((('Chondrus crispus':0.24397860919999992,'Rhodella maculata':0.10491793439999997)99:0.08284400000000003,'Galdieria sulphuraria':0.13596538999999996)92:0.04979,'Lotharella sp. CCMP622':0.6467240496)80:0.018982000000000054,'Cyanidioschyzon merolae':0.3351941785999999)92:0.037864999999999926):0.04141300000000003,(((('Anabaena variabilis':0.03880748860000005,'Nodularia spumigena':0.04779570430000002)100:0.13451600000000008,('Cyanothece sp. PCC 7425':0.051592984699999955,'Thermosynechococcus elongatus':0.12170888280000003)94:0.05369600000000008)39:0.014793999999999974,('Crocosphaera watsonii':0.12095143419999999,'Synechococcus sp. PCC 7335':0.1006569504)71:0.02967300000000006)56:0.033911000000000024,'Lyngbya sp. PCC 8106':0.10802618970000011)100:0.04141299999999992);

**Ribulose phosphate epimerase, dataset listed in Supplementary Table S18:**

((((((((((((((((((('Euglena gracilis2':0.029803000000000024,'Euglena longa1':0.09234600000000004)100:0.06878899999999999,'Nannochloropsis gaditana':0.21536599999999995)43:0.04009600000000013,('Eutreptiella gymnastica-like CCMP1594 2':0.14140200000000003,'Eutreptiella gymnastica NIES-381 2':0.07642799999999994)37:0.018836000000000075)68:0.0501459999999998,'Phytophtora ramorum':0.18699699999999986)88:0.10418600000000011,(('Cryptomonas paramecium1':0.017179999999999973,'Guillardia theta':0.26991300000000007)99:0.06569099999999994,'Rhodomonas salina1':0.21357800000000005)100:0.14700599999999997)40:0.03428000000000009,(((('Euglena gracilis1 PT':0.01215300000000008,'Eutreptiella gymnastica-like CCMP1594 1':0.10397600000000007)54:0.007885999999999838,'Euglena longa2 PT':0.05145399999999989)55:0.0455509999999999,'Eutreptiella gymnastica NIES-381 1':0.11541499999999982)100:0.14921300000000004,'Aureococcus anophagefferens1':0.455279)56:0.0193500000000002)38:0.08402200000000004,'Phaeodactylum tricornutum1':0.374231)33:0.014346999999999888,'Emiliania huxleyi2':0.3928099999999999)55:0.03425300000000009,(('Bathycoccus prasinos2':0.13975099999999996,'Micromonas pusilla2':0.35589099999999996)98:0.060462000000000016,'Ostreococcus tauri':0.18534200000000012)97:0.12658800000000014)51:0.040349999999999886,'Perkinsus marinus':0.550254)42:0.03325400000000012,('Pyramimonas amylifera1':0.2465599999999999,'Pyramimonas parkeae1':0.14924300000000001)97:0.16375400000000018)34:0.027774999999999883,'Aspergillus fumigatus':0.4531910000000001)33:0.037336000000000036,(((((('Homo sapiens':0.06359200000000009,'Xenopus laevis':0.026923999999999948)96:0.06216899999999992,'Danio rerio':0.07747900000000008)98:0.13780400000000004,('Crassostrea gigas':0.15728200000000014,'Schistosoma haematobium':0.46969700000000003)94:0.086144)67:0.03415799999999991,('Acyrthosiphon pisum':0.33798799999999996,'Drosophila melanogaster':0.14215599999999995)100:0.10403300000000004)96:0.19003900000000007,('Paramecium tetraurelia':0.5362259999999999,'Tetrahymena thermophila':0.19930000000000003)99:0.164895)62:0.06631399999999998,(('Neospora caninum':0.0444739999999999,'Toxoplasma gondii':0.08140799999999992)100:0.27813200000000005,'Picrophilus torridus':1.302514)57:0.19637799999999994)55:0.12780900000000006)48:0.08500300000000016,((((((('Entamoeba histolytica':0.5207830000000002,'Chlorella variabilis3':0.2540610000000001)59:0.10502699999999976,'Gloeochaete wittrockiana':0.2832279999999998)55:0.10712500000000014,'Acanthamoeba castellanii':0.20172000000000012)27:0.05143000000000009,((('Arabidopsis thaliana2':0.11131999999999986,'Oryza sativa2':0.08417000000000008)100:0.04288599999999998,'Selaginella moellendorffii1':0.1353740000000001)100:0.08320000000000016,'Physcomitrella patens2':0.1005060000000002)100:0.12442500000000001)24:0.022028999999999854,(((('Bigelowiella natans3':0.21081700000000003,'Lotharella sp. CCMP622':0.11680400000000013)100:0.07850899999999994,('Cryptococcus neoformans':0.42039800000000005,'Laccaria bicolor':0.18516700000000008)100:0.5229049999999997)51:0.0594260000000002,('Dictyostelium discoideum':0.27708299999999997,'Naegleria gruberi':0.39076100000000014)59:0.12712800000000013)45:0.07181099999999985,'Ectocarpus siliculosus2':0.438075)24:0.04362200000000005)47:0.04151499999999975,(('Chlamydomonas reinhardtii2':0.07479000000000013,'Volvox carteri f. nagariensis':0.14552699999999996)99:0.07933999999999974,'Dunaliella tertiolecta2':0.2659609999999999)96:0.08815199999999979)45:0.03463200000000022,'Porphyridium aerugineum1':0.3483379999999998)28:0.0284810000000002)35:0.0406979999999999,(((('Leptomonas pyrrhocoris':0.13918699999999995,'Leishmania major':0.10441699999999998)100:0.457951,'Trypanosoma brucei':0.2253989999999999)98:0.08726999999999996,'Cyanoptyche gloeocystis1':0.5676029999999999)55:0.07354499999999975,'Leishmania major2':0.8319279999999998)34:0.07672300000000032)39:0.03758000000000017,(((('Ostreococcus tauri2':0.1917629999999999,'Pyramimonas amylifera2':0.46593399999999985)73:0.10119100000000003,'Micromonas pusilla1':0.21426999999999996)100:0.19645700000000033,('Chlorella variabilis1':0.5043040000000001,'Ectocarpus siliculosus3':0.7885550000000001)92:0.10746800000000012)74:0.09797600000000006,'Cyanidioschyzon merolae':0.5555900000000003)43:0.031166999999999945)44:0.06325299999999956,'Chondrus crispus':0.3191639999999998)39:0.01948300000000014,('Corynebacter diphteriae':0.3978189999999999,'Mycobacterium tuberculosis':0.2782859999999998)100:0.24725100000000033):0.20695450000000015,((((((((((((('Bathycoccus prasinos':0.056942999999999966,'Micromonas pusilla3':0.04862999999999995)100:0.06381900000000007,'Dunaliella tertiolecta':0.11565599999999998)97:0.05014699999999994,'Pyramimonas amylifera3':0.057477)46:0.0164979999999999,('Chlorella variabilis2':0.0744689999999999,'Coccomyxa subellipsoidea':0.13423999999999991)93:0.05209099999999989)46:0.02538499999999999,('Polytomella parva':0.2497720000000001,'Pyramimonas parkeae2':0.03047399999999989)56:0.01995900000000006)65:0.030977000000000032,(((('Physcomitrella patens1':0.015273000000000092,'Physcomitrella patens3':0.014985000000000026)100:0.03815600000000008,'Arabidopsis thaliana':0.000002)100:0.021001000000000047,'Oryza sativa1':0.03148100000000009)99:0.034129000000000076,'Selaginella moellendorffii2':0.04916900000000002)99:0.04627200000000009)66:0.04443799999999998,'Chlamydomonas reinhardtii':0.1508449999999999)78:0.016858999999999957,(('Galdieria sulphuraria':0.11709199999999997,'Rhodella maculata':0.133343)88:0.07949800000000007,'Porphyridium aerugineum2':0.11243399999999992)86:0.06270599999999993)84:0.03465600000000002,'Gloeochaete wittrockiana2':0.10393399999999997)80:0.02787099999999998,((((((('Cyanothece sp. PCC 7425':0.0408980000000001,'Thermosynechococcus elongatus':0.02040799999999998)100:0.04574199999999995,'Crocosphaera watsonii':0.03262900000000002)81:0.029905999999999988,'Lyngbya sp. PCC 8106':0.012923000000000018)78:0.025060000000000082,'Synechococcus sp. PCC 7335':0.08963999999999994)93:0.027039000000000035,'Nodularia spumigena':0.06000300000000003)66:0.005044999999999966,'Anabaena variabilis':0.000002)100:0.20199499999999992,('Cyanoptyche gloeocystis2':0.000002,'Cyanoptyche gloeocystis3':0.015260999999999969)100:0.10131800000000002)90:0.06912399999999996)98:0.12207100000000004,'Cyanidioschyzon merolae2':0.16532499999999994)98:0.10902000000000012,(((((((((((((('Phaeodactylum tricornutum2':0.0503579999999999,'Thalassiosira pseudonana1':0.07356800000000008)98:0.039365000000000094,'Thalassiosira pseudonana2':0.008529999999999927)100:0.061434999999999906,'Rhodomonas salina2':0.06275300000000006)30:0.015384999999999982,'Guillardia theta2':0.07459299999999991)67:0.013965999999999923,'Cryptomonas paramecium2':0.1543460000000001)97:0.025644999999999918,(('Bigelowiella natans2':0.036243000000000025,'Lotharella sp. CCMP622 2':0.04331900000000011)70:0.015042999999999918,'Bigelowiella natans1':0.7873540000000001)88:0.07315099999999997)88:0.07494099999999992,'Aureococcus anophagefferens2':0.000002)76:0.023276000000000074,'Emiliania huxleyi':0.10868199999999995)90:0.047552999999999956,'Ectocarpus siliculosus1':0.11059400000000008)90:0.10314500000000004,'Shewanella baltica':0.07072999999999996)89:0.08025299999999991,'Burkholderia cenocepacia':0.04574999999999996)87:0.0635730000000001,'Ralstonia solanacearum':0.09305500000000011)85:0.06096699999999999,'Amphimedon queenslandica':0.30154599999999987)100:0.3367530000000001,(('Magnetospirillum magneticum':0.19289599999999996,'Nitrobacter hamburgensis':0.2931079999999999)97:0.06466400000000005,'Rhodobacter sphaeroides':0.34107599999999993)95:0.09326900000000005)62:0.061741000000000046)66:0.09133400000000003,(((('Bacteroides fragilis':0.3593709999999999,'Prevotella ruminicola':0.21726500000000004)100:0.15353299999999992,'Flavobacterium columnare':0.40093)100:0.19212300000000004,('Listeria monocytogenes':0.4910650000000001,'Staphyllococcus aureus':0.608244)64:0.09761200000000003)97:0.07321200000000005,'Listeria monocytogenes2':0.231309)95:0.09764799999999996)62:0.2069544999999997);

**Ribose-phosphate isomerase, dataset listed in Supplementary Table S19:**

(('Picrophilus torridus':0.5170400000000002,'Thermoplasma volcanium':0.45028000000000024):0.4219499999999998,(((('Acanthamoeba castellanii':0.35013000000000005,('Monosiga brevicollis':0.2967599999999999,'Gloeochaete wittrockiana3':0.2896300000000003)49:0.21989999999999998)44:0.07894999999999985,'Drosophila melanogaster':0.39154)78:0.02452999999999994,('Dictyostelium discoideum':0.4430200000000002,(('Lotharella sp. CCMP6222':0.21668999999999983,('Phytophtora ramorum':0.07853000000000021,'Pythium ultimum var. sporangiiferum':0.09296000000000015)100:0.2545099999999998)89:0.04164000000000012,('Aspergillus fumigatus':0.8693499999999998,('Cryptococcus neoformans':0.11847999999999992,'Laccaria bicolor':0.2179500000000001)100:0.35928000000000004)95:0.1942700000000004)88:0.10938999999999988)88:0.04674999999999985)100:0.3964700000000003,('Staphyllococcus aureus':1.3116699999999994,(((((('Eimeria tenella':0.7766099999999998,('Euglena gracilis 1':0.0685699999999998,'Euglena longa 3':0.08840000000000003)100:0.8842399999999997)89:0.13149999999999995,(('Cyanoptyche gloeocystis2':0.16519999999999957,'Cyanoptyche gloeocystis3':0.3183499999999997)100:0.3634900000000001,('Gloeochaete wittrockiana4':0.7320099999999998,'Gloeochaete wittrockiana':0.5073599999999998)88:0.10485999999999995)99:0.3252299999999999)61:0.01802000000000037,((('Neospora caninum':0.0014899999999999913,'Toxoplasma gondii':0.08679999999999977)100:0.39561,(((('Lotharella sp. CCMP622':0.22639999999999993,(((((('Bathycoccus prasinos':0.029380000000000184,('Ostreococcus tauri':0.05292999999999992,(((('Cryptomonas paramecium1':0.03986000000000001,('Guillardia theta':0.14525999999999994,'Rhodomonas salina':0.10926999999999998)100:0.13511000000000006)53:0.034030000000000005,'Guillardia theta2':0.04374000000000011)44:0.022930000000000117,'Cryptomonas paramecium2':0.18043999999999993)99:0.0944199999999995,((('Euglena gracilis 2 PT':0.012359999999999705,('Euglena longa 1':0.02491000000000021,'Euglena longa 2 PT':0.0)100:0.04211999999999971)76:0.01330000000000009,'Eutreptiella gymnastica NIES-381':0.10057999999999989)49:0.02245000000000008,('Eutreptiella gymnastica-like CCMP1594 1':0.0,'Eutreptiella gymnastica-like CCMP1594 2':0.0)100:0.019979999999999887)100:0.12277000000000005)87:0.01097999999999999)72:0.04611000000000054)65:0.009339999999999904,'Micromonas pusilla':0.0387000000000004)81:0.053690000000000015,('Pyramimonas amylifera1':0.09359000000000028,'Pyramimonas parkeae':0.027369999999999894)97:0.03376000000000001)78:0.03753999999999991,'Thalassiosira pseudonana2':0.011569999999999858)78:0.029290000000000038,'Thalassiosira pseudonana':0.15418999999999983)47:0.007330000000000059,'Phaeodactylum tricornutum':0.17528000000000032)62:0.017129999999999868)54:0.031179999999999986,('Emiliania huxleyi':0.20197999999999983,'Aureococcus anophagefferens':0.13804999999999978)59:0.04097000000000017)100:0.07816000000000001,((((('Chondrus crispus2':0.12809000000000026,('Galdieria sulphuraria2':0.20448000000000022,'Galdieria sulphuraria':0.08115000000000006)100:0.18116999999999983)65:0.054320000000000146,'Porphyridium aerugineum':0.06529000000000007)59:0.01134999999999975,'Rhodella maculata':0.08765999999999963)73:0.0933900000000003,'Nannochloropsis gaditana':0.12115000000000009)29:0.0,'Ectocarpus siliculosus':0.07479000000000013)73:0.05100999999999978)78:0.035719999999999974,'Cyanidioschyzon merolae':0.12212999999999985)88:0.10804999999999998)49:0.04300999999999977,(((((((('Chlamydomonas reinhardtii':0.013650000000000162,'Dunaliella tertiolecta':0.1500600000000003)77:0.011800000000000033,'Volvox carteri f. nagariensis':0.018090000000000384)78:0.02450999999999981,'Chlorella variabilis2':0.04557000000000011)63:0.02253000000000016,'Polytomella parva':0.10616000000000003)89:0.041309999999999736,('Coccomyxa subellipsoidea2':0.01831999999999967,'Coccomyxa subellipsoidea1':0.01546000000000003)100:0.08589000000000002)97:0.10637000000000008,(('Cyanoptyche gloeocystis':0.18721999999999994,'Gloeochaete wittrockiana2':0.12851999999999997)79:0.10705000000000009,((('Arabidopsis thaliana3':0.23591999999999969,'Oryza sativa3':0.07320999999999955)70:0.07705000000000028,'Selaginella moellendorffii1':0.0869500000000003)64:0.031629999999999825,('Physcomitrella patens2':0.0,'Physcomitrella patens3':0.01664000000000021)100:0.05013999999999985)64:0.06213000000000024)69:0.0384199999999999)71:0.054159999999999986,(('Arabidopsis thaliana2':0.06810000000000027,'Oryza sativa2':0.3959800000000002)97:0.07235999999999976,'Arabidopsis thaliana1':0.13293)97:0.22890999999999995)97:0.12685999999999975,'Aureococcus anophagefferens2':1.5121599999999997)17:0.0)82:0.12988000000000044)64:0.06388999999999978,((('Chlamydomonas reinhardtii2':1.5492799999999995,('Oryza sativa1':0.9206799999999999,'Physcomitrella patens1':0.3695999999999997)100:0.36064000000000007)88:0.1807000000000003,'Chlorella variabilis':1.48826)78:0.12475000000000014,'Pyramimonas amylifera2':1.2974000000000006)95:0.5643499999999997)66:0.08226000000000022,('Listeria monocytogenes':0.7982200000000002,('Nitrobacter hamburgensis':0.3365,'Rhodobacter sphaeroides':0.5265900000000001)100:0.20714000000000032)80:0.11556999999999995)56:0.11067999999999989,((('Paulinella chromatophora':0.18005000000000004,'Prochlorococcus marinus':0.16705999999999976)100:0.28309000000000006,((((((('Anabaena variabilis':0.03642999999999974,'Nodularia spumigena':0.02475000000000005)100:0.09047000000000027,'Crocosphaera watsonii':0.1705700000000001)63:0.02635999999999994,'Lyngbya sp. PCC 8106':0.09296999999999978)59:0.03716000000000008,'Synechococcus sp. PCC 7335':0.1588099999999999)96:0.10494000000000003,'Cyanothece sp. PCC 7425':0.12725000000000009)100:0.09871000000000008,'Thermosynechococcus elongatus':0.08423999999999987)100:0.1787399999999999,(('Physcomitrella patens4':0.10786000000000007,'Physcomitrella patens5':0.09074999999999989)100:0.02076000000000011,'Selaginella moellendorffii2':0.13811000000000018)100:0.40932999999999975)99:0.09065000000000012)99:0.18823999999999996,('Pyrobaculum aerophilum':0.96055,('Sulfolobus tocodaii':1.7153299999999998,((((('Burkholderia cenocepacia':0.16561000000000026,('Cupriavidus necator':0.0690900000000001,'Ralstonia solanacearum':0.10263)100:0.20176000000000016)94:0.23673999999999973,'Verminephrobacter eiseniae':0.29434000000000005)87:0.25644,'Vibrio cholerae':0.11768)59:0.05457000000000001,'Shewanella baltica':0.09065999999999974)98:0.5327999999999999,'Chondrus crispus':3.17174)33:0.03011000000000008)63:0.18101000000000012)61:0.20764000000000005)70:0.07179000000000002)57:0.09296999999999978)53:0.04167000000000032):0.02220999999999984);

**Sedoheptulose bisphosphatase, dataset listed in Supplementary Table S20:**

(((((((((((('Chlamydomonas reinhardtii':0.025648705400000082,'Volvox carteri f. nagariensis':0.013229439699999901)98:0.06462900000000005,('Chlorella variabilis2':0.1437296157000001,'Coccomyxa subellipsoidea':0.15359739109999992)64:0.026918999999999915)56:0.037806000000000006,'Dunaliella tertiolecta2':0.21121197539999992)63:0.0708390000000001,(('Euglena gracilis':0.009140148399999992,'Euglena longa':0.0564159273)100:0.19108999999999998,'Eutreptiella gymnastica NIES-381 2':0.1625442641000001)100:0.19846199999999992)57:0.03837399999999991,((('Physcomitrella patens':0.015267213500000043,'Physcomitrella patens2':0.023382119599999918)100:0.042988000000000026,'Selaginella moellendorffii':0.09452794180000001)99:0.035215999999999914,('Arabidopsis thaliana':0.04610357510000007,'Oryza sativa':0.0712128487999999)99:0.04273400000000005)100:0.1649210000000001)100:0.2118389999999999,(('Cyanoptyche gloeocystis':0.005320240399999898,'Cyanoptyche gloeocystis2':0.000002)100:0.17741099999999999,'Gloeochaete wittrockiana':0.1192653023000001)100:0.12821100000000007)93:0.14106700000000005,'Emiliania huxleyi1':1.1045736393)7:0.02599399999999985,((((('Lotharella sp. CCMP622':0.14591039449999998,'Lotharella sp. CCMP6222':0.5025148578)58:0.051415999999999906,'Ectocarpus siliculosus':0.18401767260000002)56:0.09315700000000016,(('Chondrus crispus':0.1630826824,'Porphyridium aerugineum2':0.14853726849999993)96:0.09540100000000007,'Rhodella maculata':0.13453714890000001)87:0.05380600000000002)24:0.03714700000000004,'Galdieria sulphuraria2':0.2332622577000001)46:0.08005299999999993,('Cyanidioschyzon merolae':0.4010684322999998,'Galdieria sulphuraria':0.46697512529999985)27:0.04978600000000011)11:0.0369489999999999)83:0.1394540000000002,('Cryptomonas paramecium':0.6466435332999998,'Rhodomonas salina':0.33492343319999973)96:0.13051100000000027)86:0.076241,(((((((((('Eutreptiella gymnastica-like CCMP1594 1':0.0052283186999999565,'Eutreptiella gymnastica-like CCMP1594 2':0.010484079299999838)100:0.008337000000000039,'Eutreptiella gymnastica-like CCMP1594 3':0.07310842439999998)100:0.029787000000000008,'Eutreptiella gymnastica NIES-381 1':0.1006061573999999)100:0.081596,'Guillardia theta2':0.07228276860000005)98:0.05427399999999993,'Cryptomonas paramecium2':0.11374647579999975)95:0.04022900000000007,'Guillardia theta':0.2941546532000001)99:0.11744500000000002,((('Bathycoccus prasinos':0.1522249255000001,'Ostreococcus tauri':0.07617741139999978)98:0.04032299999999989,('Micromonas pusilla':0.1212513584999999,'Pyramimonas parkeae':0.09772713519999998)98:0.04905999999999988)93:0.03894399999999987,'Phaeodactylum tricornutum':0.14191840779999954)98:0.07376100000000019)57:0.03922800000000004,('Emiliania huxleyi2':0.36921574469999996,'Emiliania huxleyi3':0.16173551979999967)99:0.14161400000000013)100:0.24824100000000016,('Eimeria tenella':0.47506808870000006,'Toxoplasma gondii':0.36583772979999996)100:0.28331099999999987)100:0.2745869999999999,('Porphyridium aerugineum':0.6276236974000002,'Rhodella maculata2':0.9478545191000001)88:0.07255300000000009)88:0.04157300000000008)82:0.1066959999999999,(((('Dunaliella tertiolecta':0.32562769829999993,'Volvox carteri f. nagariensis2':0.3419175282)100:0.21434700000000007,'Chlorella variabilis':0.45326467959999994)100:0.28153300000000003,(('Ectocarpus siliculosus2':0.2920419042,'Nannochloropsis gaditana':0.5556541062)91:0.04565700000000006,'Thalassiosira pseudonana':0.5009425613)100:0.28509000000000007)73:0.07056399999999963,('Trypanosoma brucei':0.8729297035000001,'Cyanidioschyzon merolae2':0.8205055197000002)81:0.03648999999999969)68:0.036159000000000496):0.5536830000000001,((((((('FBP_Cryptococcus neoformans':0.10029333439999988,'FBP_Laccaria bicolor':0.07212105039999983)100:0.17007400000000006,'FBP_Aspergillus fumigatus':0.4857297974999999)38:0.06260600000000016,'FBP_Emiliania huxleyi':0.7321159686)37:0.06958999999999982,'FBP_Tetrahymena thermophila':0.3503020937999999)42:0.06763800000000009,((('FBP_Chondrus crispus':0.13489096890000019,'FBP_Galdieria sulphuraria':0.15424510970000016)97:0.12614100000000006,'FBP_Dictyostelium discoideum':0.47004868899999996)97:0.06562100000000015,'FBP_Trypanosoma brucei':0.5764892642000001)43:0.08172099999999993)69:0.03588799999999992,'FBP_Arabidopsis thaliana':0.4613305413000002)92:0.11647600000000002,((('FBP_Synechococcus sp. PCC 7335':0.15430940009999983,'FBP_Thermosynechococcus elongatus':0.1802098341999998)89:0.07044499999999987,'FBP_Anabaena variabilis':0.10001159059999987)100:0.3610899999999999,'FBP_Synechococcus sp. PCC 7335_2':0.6178952746999999)98:0.291639)100:0.5536830000000001);

**Transketolase, dataset listed in Supplementary Table S21:**

(((((((((((((((((('Arabidopsis thaliana2':0.043862999999999985,'Arabidopsis thaliana1':0.04446799999999973)100:0.044877000000000056,('Oryza sativa2':0.057926000000000144,'Oryza sativa1':0.1531880000000001)100:0.026147999999999616)100:0.05328500000000025,'Selaginella moellendorffii2':0.09952200000000033)91:0.02861899999999995,('Physcomitrella patens2':0.06205999999999978,'Physcomitrella patens1':0.08019799999999977)99:0.027982000000000173)100:0.06590300000000004,('Physcomitrella patens3':0.20058299999999996,'Selaginella moellendorffii':0.24032900000000001)100:0.40313500000000024)94:0.04045999999999994,('Pyramimonas amylifera':0.0824720000000001,'Pyramimonas parkeae':0.029886999999999997)100:0.1381920000000001)92:0.03292599999999979,((((('Bathycoccus prasinos':0.10437300000000027,'Micromonas pusilla':0.07654499999999986)100:0.03238500000000011,'Ostreococcus tauri':0.33103000000000016)100:0.12325399999999975,'Coccomyxa subellipsoidea':0.1380490000000001)99:0.06339400000000017,'Chlorella variabilis':0.16334099999999996)100:0.06275899999999979,((('Chlamydomonas reinhardtii':0.0668899999999999,'Volvox carteri f. nagariensis':0.05227499999999985)100:0.05518400000000012,'Polytomella parva':0.14554899999999993)100:0.027125999999999983,'Dunaliella tertiolecta':0.1520100000000002)100:0.047787000000000024)99:0.04396299999999975)91:0.04261700000000035,(((('Chondrus crispus2':0.5833690000000002,'Chondrus crispus':0.1823640000000002)79:0.056379999999999875,'Rhodella maculata':0.18625500000000006)77:0.05995999999999979,'Porphyridium aerugineum':0.2575389999999995)62:0.05393300000000023,(('Cyanidioschyzon merolae2':0.02318700000000007,'Cyanidioschyzon merolae':0.0986290000000003)100:0.2771880000000002,'Galdieria sulphuraria':0.2784880000000003)83:0.057386999999999855)99:0.08695800000000009)84:0.03416599999999992,(('Cyanoptyche gloeocystis2':0.0229919999999999,'Cyanoptyche gloeocystis1':0.016970999999999847)100:0.1187560000000003,'Gloeochaete wittrockiana1':0.1933360000000004)100:0.08970999999999973)91:0.03433799999999998,(((((('Anabaena variabilis':0.07005599999999967,'Nodularia spumigena':0.03993000000000002)100:0.03224000000000027,'Crocosphaera watsonii':0.07004700000000019)100:0.06631900000000002,'Thermosynechococcus elongatus':0.09764700000000026)93:0.02894099999999966,(('Paulinella chromatophora':0.08408400000000027,'Prochlorococcus marinus':0.11878000000000011)100:0.1825909999999995,'Synechococcus sp. PCC 7335':0.13644799999999968)89:0.02988400000000002)84:0.013717000000000201,'Cyanothece sp. PCC 7425':0.11509299999999989)91:0.0408059999999999,'Lyngbya sp. PCC 8106':0.14715100000000003)93:0.028690000000000104)90:0.06239600000000012,((('Euglena gracilis 3 PT':0.017115999999999687,'Euglena longa 1 PT':0.0567479999999998)100:0.05257200000000006,('Eutreptiella gymnastica-like CCMP1594 1':0.05071700000000012,'Eutreptiella gymnastica NIES-381 2':0.1019779999999999)100:0.0209769999999998)100:0.05778099999999986,('Euglena gracilis 1':0.08574700000000002,'Euglena longa 3':0.06086400000000003)100:0.14220299999999986)100:0.21214400000000033)94:0.05644199999999966,(((((((('Diplonema papillatum':0.38201399999999985,'Cryptococcus neoformans2':0.7569689999999998)91:0.060203000000000007,('Trypanosoma brucei':0.2580860000000005,'Leishmania major':0.24421500000000052)100:0.20371099999999975)93:0.05155100000000035,('Naegleria gruberi1':0.3438870000000003,'Naegleria gruberi2':0.23761900000000047)100:0.35460899999999995)94:0.05940299999999965,(('Cryptococcus neoformans':0.21118799999999993,'Laccaria bicolor':0.25302099999999994)86:0.06339799999999984,'Aspergillus fumigatus':0.26766500000000004)100:0.19329600000000013)44:0.030829999999999913,(((('Perkinsus marinus3':0.06509399999999976,'Perkinsus marinus1':0.058807000000000276)96:0.050756000000000245,'Perkinsus marinus2':0.07874500000000051)100:0.29847599999999996,('Neospora caninum':0.1494620000000002,'Toxoplasma gondii':0.054088000000000136)100:0.4197009999999999)86:0.0972789999999999,(('Lotharella sp. CCMP6222':0.17334300000000002,'Lotharella sp. CCMP622':0.23589099999999963)100:0.19462800000000025,'Gloeochaete wittrockiana2':0.38269200000000003)68:0.040589999999999904)64:0.032928999999999764)55:0.02569200000000027,('Acanthamoeba castellanii':0.24649600000000005,'Percolomonas cosmopolitus':0.5010529999999997)86:0.10151900000000014)79:0.025370000000000115,((('Emiliania huxleyi2':0.5444819999999999,'Ectocarpus siliculosus2':0.33157099999999984)88:0.04140599999999983,('Phaeodactylum tricornutum2':0.31812299999999993,'Thalassiosira pseudonana':0.3141050000000001)100:0.14312499999999995)83:0.05479300000000009,'Phytophtora ramorum':0.3503750000000001)81:0.023851000000000067)89:0.0504929999999999,'Monosiga brevicollis':0.2860369999999999)100:0.1157849999999998)98:0.10208300000000037,(((((('Nitrobacter hamburgensis':0.3336039999999998,'Rhodobacter sphaeroides':0.43884199999999973)50:0.04724100000000009,'Rhodobacter sphaeroides2':0.29235299999999986)55:0.039915000000000145,'Magnetospirillum magneticum':0.29695000000000027)93:0.05107399999999984,'Burkholderia cenocepacia':0.39371599999999995)97:0.0885180000000001,'Azospirillum sp. B506':0.3747290000000003)100:0.1292089999999999,(((('Vibrio cholerae':0.09468699999999997,'Yersinia pestis':0.11237600000000025)99:0.040904000000000273,'Shewanella baltica':0.13136500000000018)100:0.13906699999999983,('Ralstonia solanacearum':0.16540600000000039,'Verminephrobacter eiseniae':0.29160299999999983)89:0.07348299999999997)90:0.045696999999999655,'Cupriavidus necator':0.3050619999999995)100:0.1527470000000002)93:0.11816100000000018)33:0.02833299999999994,((((('Bacillus anthracis':0.16498200000000018,'Listeria monocytogenes3':0.3305030000000002)86:0.04588800000000015,'Listeria monocytogenes2':0.22628400000000015)86:0.06204199999999993,'Listeria monocytogenes':0.3795010000000003)92:0.05572399999999966,'Bacillus anthracis2':0.34617799999999965)100:0.12485800000000014,('Entamoeba histolytica2':0.004107000000000305,'Entamoeba histolytica1':0.000002)100:0.5960709999999998)46:0.04212000000000016)36:0.07399500000000003,(('Mycobacterium tuberculosis':0.2448769999999998,'Streptomyces coelicolor':0.29817099999999996)92:0.05736000000000008,'Corynebacter diphteriae':0.21103300000000003)99:0.3280829999999999)78:0.12141400000000013,(((((((('Cryptomonas paramecium2':0.23068,'Guillardia theta':0.034340999999999955)80:0.023355000000000015,'Guillardia theta2':0.0442800000000001)81:0.049514999999999976,('Cryptomonas paramecium':0.15079899999999968,'Rhodomonas salina':0.05894599999999972)97:0.01232000000000033)100:0.15977700000000006,'Ectocarpus siliculosus':0.1688550000000002)97:0.04703400000000002,'Aureococcus anophagefferens':0.20622600000000002)66:0.018844000000000083,(('Phaeodactylum tricornutum':0.08674899999999974,'Thalassiosira pseudonana2':0.07185099999999967)100:0.16391800000000023,'Lotharella sp. CCMP6223':0.19770599999999972)55:0.035250000000000004)97:0.0890740000000001,'Emiliania huxleyi':0.3286070000000003)100:0.12284000000000006,((('Euglena gracilis 2':0.06586099999999995,'Euglena longa 2':0.05403400000000014)100:0.21668500000000002,('Eutreptiella gymnastica-like CCMP1594 2':0.1066600000000002,'Eutreptiella gymnastica NIES-381 1':0.20156799999999997)58:0.05191599999999985)100:0.2209080000000001,'Dictyostelium discoideum':0.24817599999999995)100:0.06470500000000001)100:0.31949099999999975)100:1.402419,((((('Homo sapiens':0.033241999999999994,'Mus musculus':0.02125200000000005)100:0.12066700000000008,'Xenopus laevis':0.11681300000000006)99:0.0866889999999998,('Bombyx mori':0.24658399999999991,'Strongylocentrotus purpuratus':0.2650100000000002)100:0.18981000000000003)100:1.0286269999999997,'Methanocaldococcus villosus':0.672304)83:0.07959000000000005,'Picrophilus torridus':0.6835649999999998)61:0.08807000000000009):0.11023249999999996,(('Sulfolobus tokodaii':0.778168,'Thermococcus sp.':0.42450199999999993)100:0.2727170000000001,'Thermoproteus uzoniensis':0.69645)78:0.11023249999999996);

**Triose-phosphate isomerase, dataset listed in Supplementary Table S22:**

((((((((((((((((((('Physcomitrella patens5':0.040072007300000045,'Physcomitrella patens6':0.048183761400000025)100:0.07879999999999998,'Physcomitrella patens4':0.04002912010000004)78:0.02761199999999997,'Physcomitrella patens3':0.07975136779999992)76:0.006335999999999897,'Physcomitrella patens':0.03346880820000009)96:0.10865299999999989,(('Arabidopsis thaliana':0.000002,'Arabidopsis thaliana3':0.000002)100:0.1077539999999999,'Selaginella moellendorffii2':0.07070510409999997)97:0.0735030000000001)96:0.03717999999999999,'Oryza sativa3':0.12602951829999998)100:0.07699600000000006,((((('Chlamydomonas reinhardtii':0.11698240589999997,'Volvox carteri f. nagariensis':0.08515219200000002)100:0.0866499999999999,'Polytomella parva':0.191855286)99:0.03646299999999991,('Dunaliella tertiolecta':0.20556350469999995,'Perkinsus marinus4':0.5401931832)99:0.15029399999999993)97:0.11990999999999996,'Chlorella variabilis2':0.3217202834999999)95:0.02635900000000002,'Coccomyxa subellipsoidea2':0.2604545669)96:0.07412200000000002)84:0.030184999999999906,(((('Entamoeba histolytica':0.43892233699999994,'Bathycoccus prasinos':0.15419648350000004)95:0.06299300000000008,('Micromonas pusilla':0.23001541600000008,'Ostreococcus tauri':0.10614411169999993)95:0.028664000000000023)96:0.09044199999999991,('Pyramimonas amylifera1':0.20381597630000003,'Pyramimonas parkeae2':0.1784122118)99:0.045366999999999935)89:0.009384000000000059,(('Micromonas pusilla2':0.14839228370000002,'Pyramimonas parkeae':0.14279933109999998)99:0.06142099999999995,'Bathycoccus prasinos2':0.21467672680000005)99:0.16389199999999993)96:0.03953999999999991)93:0.05133500000000013,((('Arabidopsis thaliana2':0.07213496639999994,'Oryza sativa':0.06448850579999998)99:0.03091099999999991,'Oryza sativa2':0.0989414340000001)99:0.03819299999999992,('Physcomitrella patens2':0.32979291359999996,'Selaginella moellendorffii':0.15010224169999997)81:0.04843299999999995)100:0.1096410000000001)77:0.048980000000000024,'Gloeochaete wittrockiana':0.38608330420000003)19:0.025167999999999857,((((((('Euglena gracilis 2':0.023077006999999927,'Euglena longa 1':0.030728591599999966)100:0.07364100000000007,('Eutreptiella gymnastica-like CCMP1594 1':0.11235014209999994,'Eutreptiella gymnastica NIES-381 1':0.07029329539999996)98:0.02032400000000001)100:0.1952360000000002,('Cyanidioschyzon merolae':0.3539966353999999,'Rhodella maculata2':0.23339265869999992)43:0.08876000000000017)42:0.026201999999999837,((('Lotharella sp. CCMP6223':0.09814002380000009,'Galdieria sulphuraria':0.3185585583000001)96:0.06917499999999999,'Rhodella maculata':0.40207256069999997)34:0.012383000000000033,('Chondrus crispus2':0.34123839590000005,'Porphyridium aerugineum2':0.1781009121999999)86:0.0819129999999999)82:0.08390000000000009)28:0.024402000000000035,(((('Euglena gracilis 4':0.026999388899999976,'Euglena longa 5':0.036761915600000084)100:0.047997999999999985,('Eutreptiella gymnastica-like CCMP1594 3':0.03674133430000004,'Eutreptiella gymnastica NIES-381 5':0.05712659860000002)99:0.04352)100:0.1037300000000001,(('Eutreptiella gymnastica NIES-381 4':0.05515630789999992,'Eutreptiella gymnastica NIES-381 6':0.0735220970999999)100:0.07766600000000001,'Eutreptiella gymnastica-like CCMP1594 4':0.067144066)100:0.352077)77:0.06706899999999982,((('Euglena gracilis 3':0.013691056999999951,'Euglena longa 3PT':0.04897482080000004)100:0.15859699999999988,'Eutreptiella gymnastica-like CCMP1594 2':0.0974367974999999)100:0.051382999999999956,'Lotharella sp. CCMP6225':0.6945126453000001)82:0.059022999999999826)62:0.04786400000000013)54:0.038270999999999944,((('Euglena gracilis 1':0.01843898910000008,'Euglena longa 2':0.030027032300000034)100:0.02230500000000002,'Euglena longa 4':0.06597307500000005)99:0.023049000000000097,'Euglena deses v. intermedia':0.05461155870000001)100:0.23648099999999994)38:0.037409000000000026,(('Acanthamoeba castellanii':0.8306158663999998,'Monosiga brevicollis':0.21121639430000005)43:0.10932199999999992,'Chondrus crispus':0.23861053359999995)18:0.013222000000000067)28:0.07179699999999989)12:0.011132000000000142,((((('Diplonema papillatum':0.43085338500000003,'Laccaria bicolor':0.23849459410000007)72:0.04758299999999993,'Aspergillus fumigatus':0.2744385118999999)62:0.03388800000000014,'Cryptococcus neoformans':0.3017969917000001)85:0.06045199999999995,'Drosophila melanogaster':0.35438827550000007)75:0.038191000000000086,'Dictyostelium discoideum':0.713396741)64:0.025442999999999882)55:0.014067000000000052,('Chlorella variabilis':0.1905983498999999,'Coccomyxa subellipsoidea':0.22783025169999993)70:0.07641900000000001)80:0.04737500000000017,(((((((((('Eutreptiella gymnastica-like CCMP1594 5':0.07752982419999999,'Eutreptiella gymnastica-like CCMP1594 6':0.06019472460000008)90:0.015066999999999942,('Eutreptiella gymnastica-like CCMP1594 7':0.06719534270000005,'Eutreptiella gymnastica NIES-381 3':0.0692696718000001)95:0.067129)99:0.03776500000000005,'Eutreptiella gymnastica NIES-381 2':0.050784182299999925)100:0.15707300000000002,'Aureococcus anophagefferens':0.20769682890000007)98:0.021999000000000102,'Emiliania huxleyi':0.17970544300000002)100:0.29368799999999995,(('Guillardia theta':0.08098621919999993,'Rhodomonas salina':0.2561899957)100:0.03996299999999997,'Cryptomonas paramecium1':0.242782799)100:0.19305600000000012)98:0.09203899999999998,'Lotharella sp. CCMP6224':0.4555364921)92:0.091148,(('Phaeodactylum tricornutum3':0.24185312790000002,'Thalassiosira pseudonana2':0.16493691519999998)100:0.1435280000000001,'Aureococcus anophagefferens2':0.3566602134000001)93:0.16279399999999988)70:0.02591600000000005,(((('Pythium ultimum var. sporangiiferum':0.08095419809999993,'Pythium ultimum var. sporangiiferum3':0.20297853310000002)71:0.061077999999999966,'Phytophtora ramorum3':0.13733091400000008)86:0.07647800000000005,'Phytophtora ramorum2':0.1149850818)100:0.2534449999999999,'Ectocarpus siliculosus2':0.33480359970000007)76:0.09481099999999998)61:0.037673999999999985,(((('Phaeodactylum tricornutum':0.13851784229999997,'Thalassiosira pseudonana':0.06845136379999994)100:0.2901959999999999,'Ectocarpus siliculosus':0.26026906289999996)39:0.08003699999999991,('Phytophtora ramorum':0.10793930809999996,'Pythium ultimum var. sporangiiferum2':0.10651469689999993)100:0.19541799999999987)93:0.10179000000000027,('Cyanoptyche gloeocystis':0.19865176880000002,'Gloeochaete wittrockiana2':0.20149564460000002)99:0.08707600000000015)66:0.03806699999999985)40:0.019559000000000104)36:0.04318399999999989,(((('Paramecium tetraurelia':0.0400755369000001,'Paramecium tetraurelia3':0.0607650663999999)100:0.18426500000000012,'Tetrahymena thermophila':0.18857421629999993)100:0.15484500000000012,'Paramecium tetraurelia2':0.48158340420000023)91:0.10022799999999998,(('Neospora caninum':0.03950816509999999,'Toxoplasma gondii2':0.012724649599999971)100:0.3850399999999998,'Nannochloropsis gaditana':0.42077524219999995)66:0.028891000000000222)65:0.04786699999999988)81:0.05366800000000005,(((('Perkinsus marinus2':0.019050079699999944,'Perkinsus marinus5':0.005523577700000004)100:0.010491000000000028,'Perkinsus marinus3':0.11278991260000004)99:0.043736999999999915,'Perkinsus marinus':0.07964381480000005)100:0.284983,('Trypanosoma brucei':0.23695351080000004,'Leishmania major':0.1359394801)100:0.306017)81:0.073013)64:0.008229000000000042,(('Lotharella sp. CCMP622':0.6393095537,'Naegleria gruberi':0.29774996259999975)90:0.09457899999999997,'Percolomonas cosmopolitus':0.5301612517999998)84:0.060199000000000114)84:0.08394899999999983,((('Neospora caninum2':0.016546345299999876,'Toxoplasma gondii':0.048956232199999894)100:0.27776899999999993,'Eimeria tenella':0.31343716919999975)100:0.08821600000000007,('Lotharella sp. CCMP6222':0.7955869855,'Porphyridium aerugineum':0.31475295589999996)95:0.07919900000000002)94:0.08510099999999987)100:0.09382050000000008,(((((((('Cryptomonas paramecium3':0.1778660475,'Phaeodactylum tricornutum2':0.5642417883000002)90:0.051026000000000016,'Cryptomonas paramecium2':0.28260288719999993)91:0.12796399999999997,'Pyramimonas amylifera2':0.40851547639999986)100:0.24795100000000003,'Cyanoptyche gloeocystis2':0.47895160429999994)68:0.18542800000000015,('Klebsiella pneumoniae':1.5013004422,'Acinetobacter johnsonii':0.7112480258)51:0.111618)68:0.160204,('Neisseria meningitidis':0.7686049125999999,'Burkholderia thailandensis':0.4484794643000001)68:0.18466000000000005)68:0.07487200000000005,(('Methylobacterium mesophilicum':0.33466249280000016,'Rhizobium tropici':0.5998879656000002)97:0.1312739999999999,'Brevundimonas diminuta':0.553110362)99:0.30996099999999993)74:0.03349000000000002,(('Escherichia coli':0.22351902779999988,'Vibrio azureus':0.2946721084999999)100:0.24208399999999997,'Pseudomonas putida':0.38148796500000004)96:0.07715300000000003):0.09382050000000008);
